# Fragile X syndrome patient-derived neurons developing in the mouse brain show *FMR1* -dependent phenotypes

**DOI:** 10.1101/2021.09.27.461739

**Authors:** Marine A. Krzisch, Hao Wu, Bingbing Yuan, Troy W. Whitfield, X. Shawn Liu, Dongdong Fu, Jennifer Shih, Carrie M. Garrett-Engele, Aaron N. Chang, Stephen Warren, Angela Cacace, Kristin R. Andrykovich, Rosalie G. J. Rietjens, Bhav Jain, Owen Wallace, Mriganka Sur, Rudolf Jaenisch

## Abstract

Abnormal neuronal development in Fragile X syndrome (FXS) is poorly understood. Data on FXS patients remain scarce and FXS animal models have failed to yield successful therapies. *In vitro* models do not fully recapitulate the morphology and function of human neurons. Here, we co-injected neural precursor cells (NPCs) from FXS patient-derived and corrected isogenic control induced pluripotent stem cells into the brain of neonatal immune-deprived mice. The transplanted cells populated the brain and a proportion differentiated into neurons and glial cells. Single-cell RNA sequencing of transplanted cells revealed upregulated excitatory synaptic transmission and neuronal differentiation pathways in FXS neurons. Immunofluorescence analyses showed accelerated maturation of FXS neurons after an initial delay. Additionally, increased percentages of Arc- and Egr1-positive FXS neurons and wider dendritic protrusions of mature FXS striatal medium spiny neurons pointed to an increase in synaptic activity and synaptic strength as compared to control. This transplantation approach provides new insights into the alterations of neuronal development in FXS by facilitating physiological development of cells in a 3D context, and could be used to test new therapeutic compounds correcting neuronal development defects in FXS.

## Introduction

Fragile X syndrome (FXS) is characterized by physical abnormalities, anxiety, intellectual disability, hyperactivity, autistic behaviors and seizures ^1^. In FXS, the expansion of the CGG triplet repeats in the *FMR1* gene leads to hypermethylation of the repeat and the *FMR1* promoter. This causes the transcriptional silencing of the *FMR1* gene after the tenth week of gestation, and the reduction or absence of *FMR1* protein (FMRP) ^2^. FMRP is an RNA-binding protein and is thought to be a translational repressor at the synapse. Previous studies have suggested that its absence causes defects in neuronal development ^1^. Although an imbalance of excitatory and inhibitory neuronal activity has been associated with seizure activity and social, communication, and behavioral challenges in several Autism Spectrum Disorders (ASDs) as well as in corresponding animal models ^3–5^, data on FXS patients are scarce. Morphologically, increased spine density and increased numbers of long and immature-appearing spines were found in the cortices of FXS patients, highlighting defects in neuronal connectivity in FXS ^6^.

So far, findings from animal models of FXS have failed to translate into successful therapies, highlighting the need to develop human cell-based models of FXS. Several studies have used neurons derived from human induced pluripotent stem cells (iPSC) or human embryonic stem cells (ESC) cultured *in vitro* ^7–10^, with some studies reporting impaired maturation and synaptic hypoactivity of FXS neurons ^7, 8, 10, 11^, and other studies showing synaptic hyperactivity and faster acquisition of synapses in FXS neurons ^12–14^. Canonical two-dimensional culture conditions failed to fully recapitulate the morphological and functional characteristics of neurons in the human brain and showed significant intrinsic variation. Additionally, a recent study showed that stress pathways were ectopically activated in cerebral organoids, the most commonly used 3-dimensional *in vitro* model, impairing cell type specification. These defects were alleviated by transplantation of the organoids in the mouse brain cortex ^15^. This finding suggests that transplantation of human neural progenitor cells (hNPCs) into the mouse brain may better reflect cellular behavior in the human brain than growth in the culture dish. Previous studies have shown that once transplanted into the neonatal brain, hNPCs migrate away from the injection site, differentiate into neurons, undergo maturation, express brain region-specific markers and become electrically active ^16–18^.

Here, we used co-transplantation of hNPCs differentiated from FXS patient-derived iPSC and isogenic control iPSC in the brain of immune-deprived mouse neonates to study developmental defects of FXS neurons. We assessed the effects of *FMR1* silencing on the development of human neurons in an *in vivo* context. Gene list functional enrichment analysis of single cell RNA sequencing data from transplanted FXS and isogenic control neurons showed upregulation of pathways linked to neurogenesis, neuronal differentiation and synaptic signaling in FXS neurons. Transplanted FXS neurons displayed an initial maturation delay, followed by accelerated maturation as compared to isogenic control, and an increased percentage of FXS neurons was positive for Arc and Egr1, two markers of neuronal activity. Additionally, FXS striatal medium spiny neurons (MSNs) had wider dendritic synaptic protrusions at 6-7 months post-injection (PI). Our data indicate that FXS neurons transplanted in the mouse brain show accelerated maturation after an initial delay, and suggest that their synapses are hyperactive as compared to isogenic controls.

## Material and Methods

### Experimental Animals

The animals used for this study were NOD/SCID/gamma mice. They were kept in group housing under standard barrier, light and temperature-controlled conditions. Food and water were available *ad libitum*. Every effort was made to minimize the number of animals used and their suffering, and all experiments were performed in accordance with the Department of Comparative Medicine and Massachusetts Institute of Technology animal husbandry standards.

### Induced pluripotent stem cell (iPSC) culture and Neural progenitor cell (NPC) differentiation

FXS patient-derived iPSC lines and isogenic control cell lines used in this study are listed in Table S3. iPSCs were cultured in feeder-free conditions on Geltrex (ThermoFisher Scientific, A1413302) or Matrigel coated flasks in StemFlexTM medium (ThermoFisher Scientific, A3349401). Cells were passaged using ReLeSR^TM^ (STEMCELL Technology, 05873) and tested for mycoplasma contamination and karyotypic abnormalities.

NPCs were induced with dual SMAD inhibition using PSC Neural Induction medium (ThermoFisher Scientific, A1647801) according to the manufacturer’s protocol. Briefly, when reaching ∼80% confluency, iPSCs were dissociated with Accutase^TM^ Cell Dissociation Reagent (ThermoFisher Scientific, A1110501) at 37°C for 5 min. Cell pellets were resuspended in medium after centrifugation and cell density was measured. To induce neural precursor cell (NPC) differentiation, single cells were plated at 2.5 × 10^5^ /cm^2^ in StemFlex^TM^ medium with 10 µM rock inhibitor (Stemgent Y27632). 48 hours after plating, medium was changed to PSC Neural Induction medium. The cells were fed every day. 6 days after induction, immature NPCs were purified with MACS sorting using anti-PSA/NCAM microbeads (Miltenyi Biotec, 130-092-966). NPCs were further expanded with neural progenitor cell expansion medium. For maintenance and further expansion, NPCs were cultured in STEMdiff™ Neural Progenitor Medium (STEMCELL Technologies, 05833) for up to 8-10 passages.

### Lentiviral labeling of NPC

Lentiviruses carrying the expression cassette of GFP or mCherry were produced by transfecting HEK293T cells with FUW constructs together with standard packaging vectors (pCMV-dR8.74 and pCMV-VSVG) followed by ultra-centrifugation-based concentration. NPCs were infected with these viruses. Once strong expression of GFP or mCherry was visible, labeled NPC were purified by FACS sorting and amplified on Geltrex or Matrigel using STEMdiff™ Neural Progenitor Medium.

### NPC transplantation

Cultured NPCs were dissociated using Accutase and resuspended in phosphate buffer saline without calcium and magnesium prior to injection, at a concentration of 10^5^ cells/μL. Post-natal day 0 to post-natal day 3 mouse pups of either gender were manually injected with a total of 4×10^5^ NPC dispersed over four injection sites in the lateral ventricles (two injection sites per brain hemisphere, one anterior and one posterior) using glass micropipettes.

### Transplanted human cell extraction for single-cell RNA sequencing

Mice were euthanized using cervical dislocation, and the brains were extracted and dissociated using Miltenyi Adult mouse and rat Brain Dissociation Kit and the gentleMACS Octo Dissociator (Miltenyi Biotec 130-107-677). The cells were then resuspended in PBS + 0.5% BSA and stained with DAPI to determine viability. DAPI-negative GFP-positive and DAPI-negative mCherry-positive cells were isolated using FACS sorting and immediately sent for single-cell RNA sequencing. Between 10^4^ and 10^5^ cells were obtained for each brain for each group (FXS and isogenic control).

### Single-cell RNA sequencing and data analysis

The cells were sequenced immediately after extraction and FACS sorting. 5000 cells were targeted. Sequencing data were mapped to a reference meta-genome composed of human hg38 (GRCh38), mouse mm10 (GRCm38), GFP and mCherry sequences. The reference genome index was created using STAR^19^ (v.2.5.1b) from Cell Ranger (v.3.1.0) with the default parameters for cellranger mkref and the following settings: --alignIntronMax 6000000 --sjdbOverhang 59. We used the Ensembl version 97 gene annotation to assign UMIs to genes. Contamination of the samples by murine cells was very small, as only 0.89%, 2.66%, 5.94% and 3.60% of total cells had more than 70% of UMIs mapped to mouse genes in control 1, control 2, FXS 1, and FXS 2 respectively. Only the cells with more than 95% of UMIs mapped to human genes were used for subsequent analysis.

The inverted beta-binomial test was used to compare paired count data between FXS and control samples with the countdata^20^ package for R^21^. The cell counts were normalized so that each sample had the same total counts and the Benjamini-Hochberg correction^22^ was applied for multiple hypothesis testing.

#### Single-cell RNA-seq analysis

Seurat v3 ^23^ was used for quality control and analysis of the single cell RNA-seq experiment. Following the Seurat Guided Clustering Tutorial (https://satijalab.org/seurat/pbmc3k_tutorial.html), cells of low quality or with too few reads were removed. The following cut offs were applied: “nFeature_RNA” > 500, “nCount_RNA” < 40000 and “percent.mt” < 20. Because differences in cell cycle among individual cells can be an unwanted source of heterogeneity, these effects were mitigated by regressing out the difference between the G2/M and S phase scores, where the scores were based upon canonical markers of cell cycle. Cells were embedded in a k-nearest neighbors (kNN) graph based on the Euclidean distance in the space of the 30 leading principal components. The quality of the cell partitions to clusters was optimized by applying the Louvain algorithm with a resolution of 0.2.

#### Differential Gene expression in scRNA-seq

The muscat^24^ (Multi-sample multi-group scRNA-seq analysis tools) package for R^21^, which can accommodate experimental designs like ours with paired replicates, was used to detect differential gene expression. After the processing steps described above, muscat was used to group cells based on metadata such as samples, clusters and conditions and tabulate pseudo-bulk RNA-seq counts by taking the sum of raw single cell counts. Differential state analysis was carried out using edgeR^25^ with a design matrix for paired samples. Genes with locally adjusted p-values less than 0.05 and at least two-fold effect sizes were considered to be differentially expressed (DE) here.

#### Gene list enrichment analysis

We performed gene list enrichment analysis on the DE protein-coding genes between control and FXS using ToppFun from the ToppGene suite ^26, 27^. The probability density function method was used to estimate p-values and a pathway significance threshold of FDR<0.05 was set. The Benjamini-Hochberg procedure^22^ was used to correct for multiple hypothesis testing.

#### Gene Set Enrichment Analysis (GSEA)

To check if cell death was different between the FXS and control, we performed GSEA^28^ on the apoptosis gene-set from the Hallmark collection of the MSigDB^28^ using log_2_ fold changes to pre-rank genes. To mitigate the influence of noise from weakly expressed genes (Suppl. Figure 1) in this analysis, genes that were expressed in less than 1% of cells were excluded.

#### Trajectory analysis

We used principal component analysis (PCA) to generate pseudotime inference with Slingshot ^29^. The ordering of clusters along the pseudotime coordinate was consistent, regardless with clusters assigned using Seurat or treating the data as a single cluster. Two lineages were identified with NPC 1 as a starting point. The first lineage led from NPCs to immature neurons to more mature neurons. The second lineage was from NPC 1 to glial cells. In addition, we used the destiny^30^ (v.3.0.1) package for R^21^ to draw diffusion maps to define differentiation trajectories.

### DNA Methylation analysis

Pyro-seq of all bisulfite-converted genomic DNA samples was performed with the PyroMark Q48 Autoprep (QIAGEN) according to the manufacturer’s instructions. The primers for pyro-seq of the *FMR1* promoter are listed below.

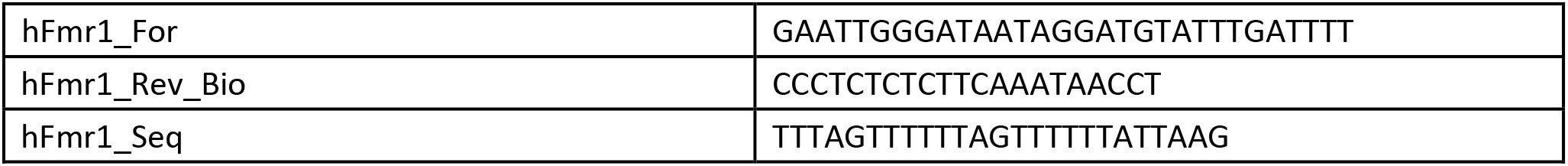

### Immunohistochemistry and immunocytochemistry

Cells in culture were fixed using 4% PFA. For the IHC analysis of cells transplanted in the mouse brain, mice at 1, 3 or 6 months post-transplantation were perfused with 4% FPA through transcardial route. 100 µm-thick brain sagittal slices were sliced with Cryostat tissue slice instrument (Leica) after cryopreservation in 30% sucrose. Immunostainings were performed using antibodies listed in Table S4. For analyses of Arc expression by IHC, a rabbit anti-FMRP antibody (Cell Signaling Technology 7104) was labeled with Alexa fluor 555 fluorophore using an antibody labeling kit (Thermo Fisher Scientific A20187). Images were acquired using a Zeiss LSM700 confocal microscope. Antibodies used in this study are listed in Table S2.

### Neuronal imaging, tracing and morphometric analysis

Cells were identified as striatal medium spiny neurons using colocalization of DARPP32 immunofluorescence staining with the cell body.

#### Neuronal arborization analysis

Confocal microscopy z-stacks of control and FXS neurons were acquired with a 20x dry objective with zoom 0.6 using a Zeiss LSM 700 confocal microscope. Confocal microscopy z-stacks of neurons were converted to grayscale images with Fiji software (ref) and traced using Neuromantic neuronal tracing freeware (Darren Myat, http://www.reading.ac.uk/neuromantic) in semi-automatic mode. After tracing, Neuromantic generated a vector file recording morphometric data including the number of bifurcations, stems, cables (branches) and terminals. The vector files were exported to Excel for data analysis.

#### Dendritic spine width and density analysis

Confocal microscopy z-stacks of control and FXS neurons were acquired with a 40x oil objective with zoom 4 using a Zeiss LSM 700 confocal microscope. The z-stepsize was chosen so that two contiguous confocal plans overlap by 50%. Images were deconvolved using Huygens and a theoretical point-spread function. 3-dimension automated analysis of dendritic protrusion density and width was performed using the Filament Tracer function of the Imaris software (RRID:SCR_007370) using a previously published protocol (Swanger et al Molecular Brain 2011). The maximum spine length and minimum spine end diameter were set at 5 μm and 0.215 μm

#### Statistical analyses of immunofluorescence experiments

Statistical analyses were performed using GraphPad Prism (v. 8) software (GraphPad software, San Diego, California, USA, www.graphpad.com). Error bars in the figures indicate standard error of the mean (SEM). For neuronal maturation analysis, we performed a two-way repeated measures ANOVA test followed by a Šídák correction for multiple comparisons. For Arc expression analysis, we performed a paired t-test to compare the percentages of Arc+ FXS and isogenic control neurons. For dendritic spine protrusion tip diameter and density analysis, Shapiro–Wilk tests were performed on each group of data to test for distribution normality. When the distribution was not normal, the non-parametric Mann–Whitney test was applied. When the distribution was normal, the equality of variances of the groups was tested and an unpaired t-test was used. For the analysis of the percentage of small, medium and large protrusions and the analysis of the number of bifurcations, cables, stems and terminals per neuron, a mixed-effects model was used, followed by a Šídák’s multiple comparisons test.

## Results

### Differentiation and migration of hNPCs transplanted into the neonatal brain

To investigate the developmental phenotypes of FXS neurons *in vivo*, we used two different FXS/corrected isogenic control pairs. In the FXS_SW/ C1_2_SW pair, the isogenic control C1_2_SW was generated by CRISPR-mediated deletion of the CGG repeats in the FXS patient-derived male FXS_SW iPSC line ^31^, leading to the reactivation of the *FMR1* promoter and FMRP re-expression (deletion pair). In the dCas9-Tet1/dCas9-dTet1 pair, the FXS patient-derived male FXS2 iPSC line was targeted with a catalytically inactive Cas9 fused to Tet1, a methylcytocine dioxygenase, as previously described ^12, 32^. This led to the demethylation of the CGG repeats, the reactivation of the *FMR1* promoter and FMRP reexpression (dCas9-Tet1 iPSC line). To generate a control where the CGG repeats were not demethylated, the FXS2 iPSC line was targeted with a catalytically inactive Cas9 fused to a catalytically inactive Tet1 (dCas9-dTet1 iPSC line) (Suppl. Figure 2). To eliminate the potential off-target effects associated with methylation editing in our phenotypical analysis, we compared the dCas9-dTet1 iPSC line, which was expected to display an FXS phenotype, and its isogenic control dCas9-Tet1 iPSC line (demethylation pair) in all the experiments. Both the demethylation and deletion of the CGG repeats rescued FXS neuronal activity phenotypes *in vitro* ^12, 13^.

hNPCs were generated from mutant and isogenic control iPSCs and sorted for PSA-NCAM expression using magnetic-activated cell sorting (MACS) in order to isolate hNPCs directed towards a neuronal fate. FXS and control NPCs were labeled with different reporters (GFP or mCherry) in order to distinguish FXS cells from control cells after transplantation into the mouse brain (Figure 1A). We characterized the hNPCs using immunofluorescence. Most cells were Sox2-Pax6-and Nestin-positive: 89%±4 C1_2_SW GFP hNPC, 87%±0.4 FXS_SW mCherry hNPC, 91%±0.04 dCas9-Tet1 and 75%±7 dCas9-dTet1 hNPC co-expressed Pax6, Sox2 and Nestin, three hNPC markers, showing high purity of cultured hNPCs. 10%±8 dCas9-dTet1 NPCs were only positive for Sox2 and Nestin (Suppl. Figure 3). In the following, the FXS_SW/ C1_2_SW and dCas9-Tet1/dCas9-dTet1 pairs will be referred to as deletion pair: control (deletion)/FXS and as demethylation pair: control (demethylation)/FXS, respectively.

**Figure 1:**
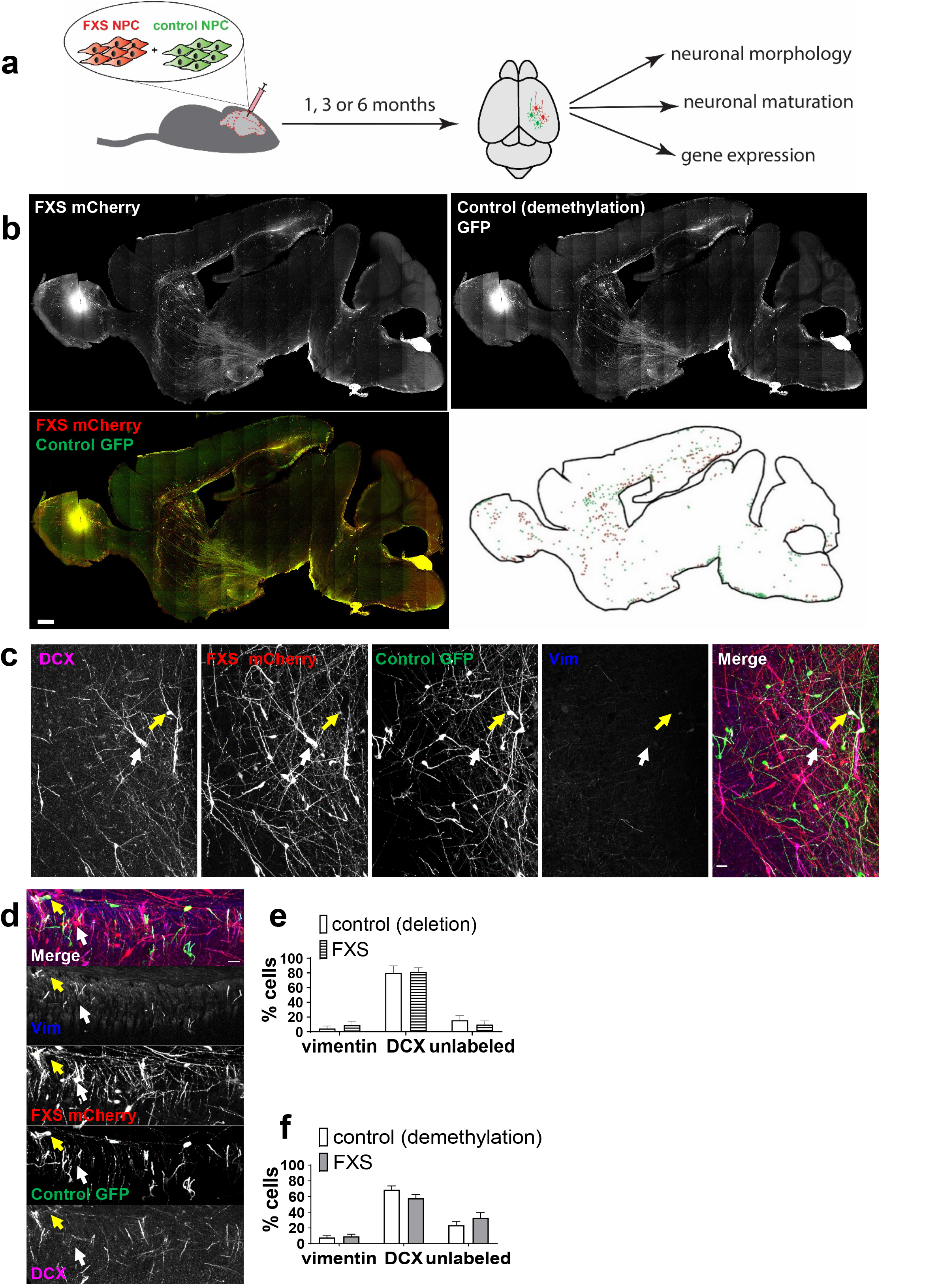
Transplanted hNPCs populate several areas of the brain and differentiate into neurons and astrocytes. **a.** Co-injection of FXS hNPCs labeled with mCherry and isogenic control hNPCs labeled with GFP in the cerebral ventricles of NSG mouse neonates. Brains were harvested at 1, 3 or 6 months post-injection (PI). **b.** Maximum intensity projection of a 100 µm-thick mouse brain slice at 1 month PI. Scale bar represents 200 µm. Red: mCherry-labeled FXS; green: GFP-labeled isogenic control (demethylation). **Right lower panel:** localization of the cell bodies of transplanted FXS (red) and isogenic control (green) cells. **c. Upper panel:** Transplanted hNPCs from FXS and isogenic control (demethylation) labeled with doublecortin (DCX) at 1 month PI. **Lower panel:** Transplanted FXS and isogenic control (demethylation) hNPCs labeled with vimentin (vim) at 1 month PI. Red: mCherry-labeled FXS; green: GFP-labeled isogenic control. The Yellow and the white arrow show the cell body of a control and FXS cell respectively. Scale bars represent 50 µm. **d.** Percentage of FXS and isogenic control (deletion) cells labeled with vim, DCX, or unlabeled at 1 month PI. 2-way repeated measures ANOVA; N= 4 animals, 17 to 109 neurons per animal for each group. **e.** Percentage of FXS and isogenic control (demethylation) cells labeled with vim, DCX, or unlabeled at 1 month PI. 2-way repeated measures ANOVA; N= 4 animals, 118 to 140 neurons per animal for each group. Bar heights and whiskers represent the mean +/-SEM.

We co-injected hNPCs derived from FXS and isogenic control cell lines into the brain ventricles of immune-deficient mouse neonates (Figure 1A) and analyzed the brains at different time points after injection. Transplanted hNPCs migrated through the mouse brain and populated several areas including the hippocampus, cortex, striatum, thalamus, and midbrain (Figure 1B). Co-staining for vimentin and doublecortin showed that most of the hNPCs had differentiated into doublecortin (DCX)-positive immature neurons at 1 month post-injection (PI) with a small proportion of hNPCs (∼10%) being positive for vimentin (vim), a marker of glial cells and neural progenitor cells (NPCs) (Figure 1C, D, E). A small proportion of cells expressed Olig2, a marker of the oligodendrocyte lineage expressed in all cells of the oligodendrocyte lineage^33, 34^, in the deletion pair (suppl. Figure 4): 7±3% control and 0.9±0.9% FXS (4 animals, 101 to 138 cells analyzed per animal). No transplanted cell was expressing Olig2 at 1 month PI for the demethylation pair (4 animals, 100 cells analyzed per animal). Importantly, oligodendrocyte progenitors can also express doublecortin^35^, although at lower levels than immature neurons. We therefore performed doublecortin and Olig2 co-staining on the mouse brain slices at 1 month PI and found that transplanted cells did not co-stain for Olig2 and doublecortin (suppl. Figure 3). Additionally, no vimentin- and doublecortin-staining was found (Figure 1C, D, E). This indicates that our antibody against doublecortin can be used to reliably detect immature neurons at 1 month PI. FXS and isogenic control hNPCs yielded similar fractions of immature neurons at 1 month PI (Figure 1C-E). No obvious difference in migration was detected between FXS and isogenic control. All the transplanted cells expressed human nuclei antigen as assessed by immunofluorescence, confirming their human origin (suppl. Figure 5). ∼60 to 80% of dCas9-Tet1 (control, demethylation pair) and C1_2_SW (control, deletion pair) transplanted neurons were FMRP-positive at 1, 3 and 6 months PI. In contrast, none of the FXS_SW (FXS, deletion pair) and FXS2 (FXS, demethylation pair) cells were FMRP-positive at the same time points (Suppl. Figure 5A,B,C). This shows that transplanted isogenic control neurons retain stable FMRP expression *in vivo* over time. Moreover, the demethylation of the *FMR1* promoter was maintained in the epigenetically edited isogenic pair, and RT-qPCR of total mRNA showed robust expression of the *FMR1* gene in C1_2_SW (control, deletion pair) cells at 1 month PI (suppl. Figure 6D,E,F).

### Altered maturation process of transplanted FXS neurons

DAPI is predominantly impermeable to live cells and can be used as a cell viability dye, DAPI-negative cells being considered viable. In order to investigate gene expression phenotypes of transplanted FXS neurons, we collected GFP+DAPI- and mCherry+DAPI-isogenic control and FXS viable transplanted neurons from the deletion pair by whole-brain extraction and subsequent FACS sorting and performed single-cell RNA sequencing on the cells using the 10x Genomics platform. We analyzed two engrafted mouse brains at 1 month PI.

FXS and isogenic control cells were assigned to similar UMAP clusters, allowing us to compare FXS and isogenic cells within the same clusters (Figure 2A). To identify the cell types corresponding to the different clusters, we used canonical marker genes, cell cycle markers and a comparison with a published single cell RNA seq data from human embryonic primary cortex^36^ (Figure 2B, C, suppl. Figure 7). The cells did not express markers specific for microglia (CXCR1… ITGAM) or oligodendrocyte progenitor cells (CSPG4…PDGFRA) (Figure 2C). Two clusters were identified as immature neurons, as they did not express NPC markers (CXCR4…REST) (Figure 2C), were not proliferative (Suppl. Figure 7B) and expressed neuronal markers (ENO2…RBFOX3). Two clusters were identified as more mature neurons, although they were still expressing DCX and therefore were still immature, as is expected at this time point after injection (Suppl. Figure 7A). These clusters expressed neuronal markers at higher levels than the two immature neuronal clusters (Figure 2C, Suppl. Fig 7C). Two clusters were identified as neural progenitor cells (NPC) via expression of markers for NPC and proliferation (Figure 2C, Suppl. Figure 7B). One cluster was named Glial cells as it expressed markers for astrocytes (GFAP… NES). Within the glial cell cluster, a subpopulation of cells expressed OLIG1 and OLIG2, markers of oligodendrocytes (Figure 2C). However, OLIG1 and OLIG2 can also be expressed by astrocytes and the cells were not expressing other markers of oligodendrocytes or oligodendrocyte progenitors such as OLIG3, MOG and CLDN11 (Figure 2C), therefore we did not define this subcluster as oligodendrocytes. To confirm these cell type assignments, we compared our samples with those from a published scRNA seq study of primary human neocortex^36^. The expression patterns of cell type marker genes in the transplanted neurons recapitulated those of similar cell types from primary cortex (Figure 2B, C). For example, the transplanted Glial cells cluster had an expression pattern that was similar with vRG (ventricular radial glia) from the primary reference. Likewise, the transplanted Neural progenitors 1 cells had a similar expression pattern with the PgG2M and PgS (cycling progenitors in G2/M phase and in S phase, respectively), and transplanted Immature neurons 1 had a similar expression pattern with ExM (Maturing excitatory neurons). To obtain an unbiased characterization of cellular dynamic processes, we used Slingshot ^29^ to assign a pseudotime to each cell, representing where the cell is along developmental trajectories. The trajectory analysis showed that cells transitioned from NPCs (clusters NPC 1 and NPC 2) to immature neurons (Immature neurons 1 and Immature neurons 2) and then to more mature neurons (cluster More mature neurons 1 and More mature neurons 2) (Suppl. Figure 7D, E). Together, these results show that the cells collected at 1 month PI were composed of neurons and NPCs at different maturation stages, and glial cells.

**Figure 2:**
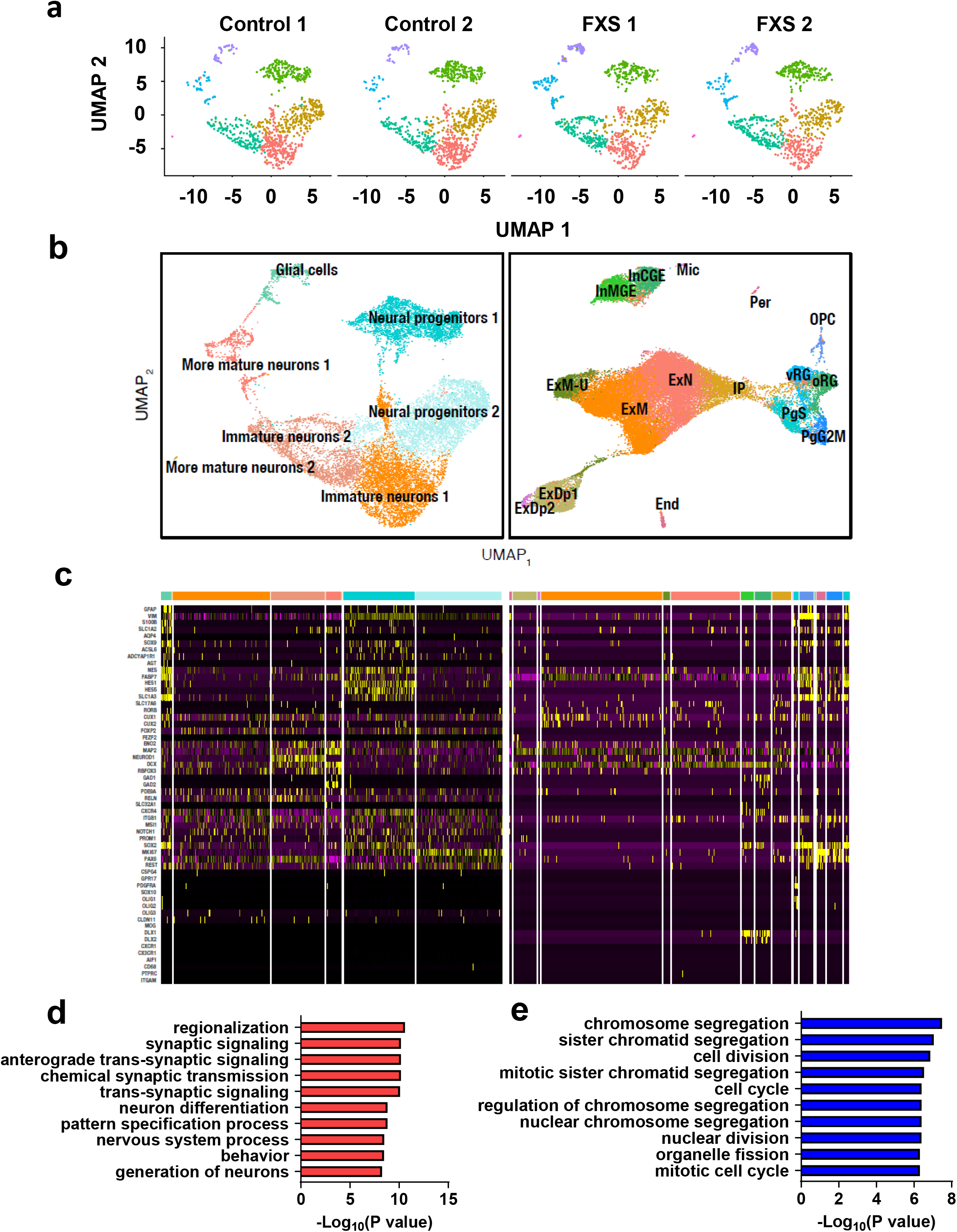
Single cell RNA sequencing of transplanted isogenic control and FXS neurons suggests increased maturation of FXS neurons and reveals upregulation of synaptic transmission, neuronal differentiation and neuronal development pathways in FXS neurons. **a.** Separate UMAP clusters for isogenic control and FXS cells from the deletion pair. 1000 cells were randomly selected from each sample. **b.** UMAP embeddings of single cell RNA-seq data are displayed for human-mouse chimeras (left panel) and mid-gestation primary human neocortex (right panel) from a published study^36^. The primary cells include outer and ventricular radial glia (oRG, vRG), G2/M and S phase cycling progenitor cells (PgG2M, PgS), intermediate progenitor cells (IP), migrating excitatory neurons (ExN), migrating excitatory upper enriched neurons (ExM-U), maturing excitatory neurons (ExM), excitatory deep layer neurons (ExDp1, ExDp2), interneurons (InCGE, InMGE), oligodendrocyte progenitor cells (OPC), endothelial cells (End), pericytes (Per) and microglia (Mic). **c.** Marker genes identify cell types in transplanted human cell model and primary neocortex. Canonical marker genes for different cell types, including astrocytes (GFAP… NES), excitatory neurons (FABP7…FEZF2), immature neurons (ENO2…RBFOX3), inhibitory neurons (GAD1, GAD2…SLC32A1), NPCs (CXCR4…REST), OPCs (CSPG4…PDGFRA), oligodendrocytes (SOX10… CLDN11) and microglia (CXCR1… ITGAM) were used to make heatmaps of gene expression for the transplanted cells (left panel) and primary neocortex (right panel). For the transplanted cells, the cluster order is from left to right: Glial cells, Immature neurons 1, Immature neurons 2, More mature neurons 1, More mature neurons 2, Neural precursors 1, Neural precursors 2. For the primary human cortex, the order of the clusters is, from left to right: Endothelial cells (End), Excitatory deep layer neurons 1 (ExDp1), Excitatory deep layer neurons 1 (ExDp2), Maturing excitatory neurons (ExM), Migrating excitatory upper enriched neurons (ExM-U), Migrating excitatory neurons (ExN), Interneurons 1 (InMGE), Interneurons 2 (InCGE), intermediate progenitor cells (IP), Microglia (Mic), Oligodendrocyte progenitor cells (OPC), Outer radial glia (oRG), Pericytes (Per), G2/M phase cycling progenitor cells (PgG2M), S phase cycling progenitor cells (PgS), Ventral radial glia (vRG). **d.** Ten most significantly upregulated gene pathways in FXS neurons. **e.** Ten most significantly downregulated gene pathways in FXS neurons. Up- or downregulation of pathways was considered significant when FDR<0.05.

FMR1 was significantly upregulated in control neurons compared to FXS, however, only 7% of control cells showed FMR1 expression (Suppl. Figure 8). This is likely due to the low sequencing depth of the 10x genomics platform and/or the low level of FMR1 expression in the cells as 60% of FXS neurons were positive for the FMR1 protein, Fmrp, as assessed by immunofluorescence. The most strongly upregulated genes in FXS neurons included a known target of Fmrp, NKX2-2^37^. Additionally, the most strongly downregulated genes in FXS neurons included HES5, which is downregulated by two putative targets of Fmrp, miR-219 and miR-338^38^. This is consistent with the role of Fmrp as a translational repressor (Tables S1 and S2). Gene list enrichment analysis of the differentially expressed genes between FXS and isogenic neuronal clusters (Immature neurons 1, Immature neurons 2, More mature neurons 1, More mature neurons 2) indicated the upregulation of neurogenesis, neuronal differentiation and synaptic signaling (Figure 2C). During neuronal differentiation, neural progenitors exit the cell cycle and differentiate into neuroblasts. Gene list enrichment analysis of the differentially expressed genes in the neuronal clusters revealed downregulation of pathways involved in cell division (Figure 2D). We analyzed the numbers of FXS and isogenic control cells in each cluster, and found that the distribution of the FXS cells was shifted towards more advanced neuronal differentiation stages: the proportion of NPCs in FXS cells was decreased, whereas the proportion of immature and mature neurons was increased (suppl. Figure 9). As this may be due to increased cell death of control neurons compared to FXS neurons after brain extraction and FACS sorting, we performed gene set enrichment analysis (GSEA)^28^ on the single cell RNA seq data from the control and FXS samples. No significant upregulation of the apoptosis pathway could be found in FXS NPCs and neurons (Suppl. Figure 9B, C). Additionally, the percentage of mitochondrial genes was similar between FXS and control cells (Suppl. Figure 9D). This suggests that there is not difference in cell death between control and FXS cells after extraction and FACS sorting. Together, these results suggest increased maturation of FXS cells at 1 month PI.

In order to further investigate alterations in the maturation and synaptic activity of FXS neurons, we analyzed transplanted neurons on brain slices using immunofluorescence. We investigated whether FXS neurons showed increased or accelerated maturation compared to isogenic controls using immunofluorescence on brain slices from transplanted mice. Neuroblasts initially express doublecortin (DCX), an immature neuron marker and gradually lose DCX expression and start expressing NeuN, a mature neuron marker ^39^. The proportion of DCX-positive neurons decreased and the proportion of NeuN-positive neurons increased between 15 days PI and 3 months PI (Figure 3 A-G), showing ongoing neuronal maturation. At 6 months PI, no DCX-positive cell could be found, suggesting that transplanted neurons had undergone full maturation.

**Figure 3:**
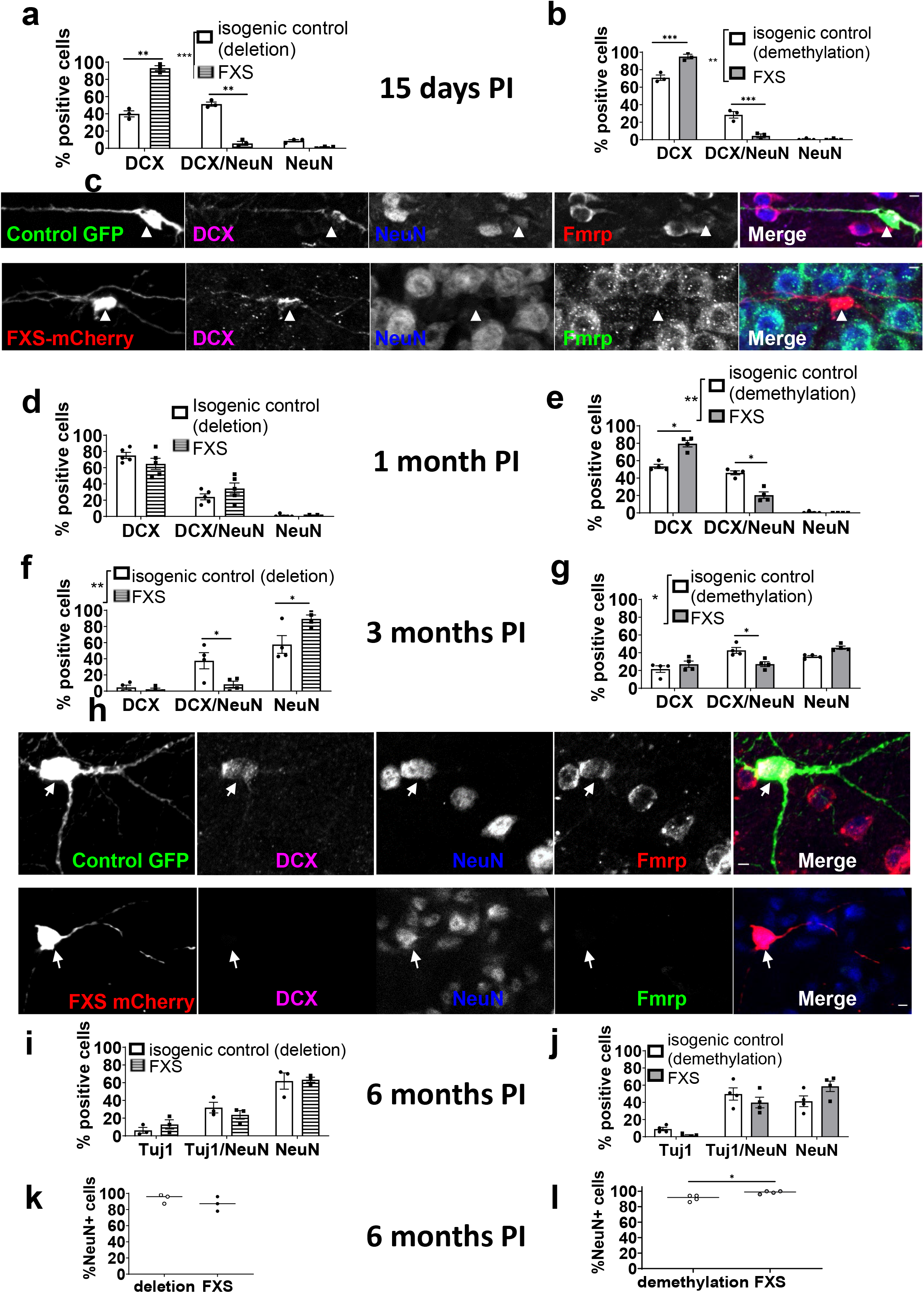
Altered maturation process of transplanted FXS neurons compared to isogenic control. Percentage of doublecortin+ (DCX), doublecortin+ NeuN+ (DCX/NeuN) and NeuN+ (NeuN) cells at 15 days post-injection (PI) in control and FXS cells from the deletion pair (**a**) and from the demethylation pair (**b**). Repeated measures two-way ANOVA followed by Šídák’s multiple comparisons test; N=3 mice, 51 to 55 neurons per mouse per group. **c.** Confocal maximum intensity projection of isogenic control (deletion) (**upper panel**) and FXS (**lower panel**) neurons immunostained with doublecortin (DCX) and NeuN at 15 days PI. White arrows indicate the cell bodies of neurons. The FXS neuron is stained with DCX only, whereas the control neuron shows DCX/NeuN co-staining, indicating a more advanced stage of maturation of the control neuron. Scale bars represent 5 µm. Percentage of doublecortin+ (DCX), doublecortin+ NeuN+ (DCX/NeuN) and NeuN+ (NeuN) cells at 1 month post-injection (PI) in control and FXS cells from the deletion pair (**d**) and from the demethylation pair (**e**). Repeated measures two-way ANOVA followed by Šídák’s multiple comparisons test; N=4 mice, 30 to 72 neurons per mouse per group. Percentage of doublecortin+ (DCX), doublecortin+ NeuN+ (DCX/NeuN) and NeuN+ (NeuN) cells at 3 months PI in control and FXS cells from the deletion pair (**f**) and from the demethylation pair (**g**). Repeated measures two-way ANOVA followed by Šídák’s multiple comparisons test; N=4 mice, 45 to 73 neurons per mouse per group. **h.** Confocal maximum intensity projection of isogenic control (demethylation) (**upper panel**) and FXS (**lower panel**) neurons immunostained with doublecortin (DCX) and NeuN at 3 months PI. The FXS neuron is stained with NeuN only, whereas the control neuron shows DCX/NeuN co-staining, indicating a more advanced stage of maturation of the FXS neuron. White arrows indicate the cell bodies of neurons. Scale bars represent 5 µm. Percentage of Tuj1+, Tuj1+ NeuN+ (Tuj1/NeuN) and NeuN+ (NeuN) cells at 6 months PI in control and FXS cells from the deletion pair (**i**) and from the demethylation pair (**j**). Repeated measures two-way ANOVA; N=3 to 4 mice, 49 to 55 neurons per mouse per group. **i.** Total percentage of NeuN+ cells at 6 months PI in control and FXS cells from the deletion pair (**k**) and from the demethylation pair (**l**). Paired t-test; N=3 to 4 mice, 49 to 55 neurons per mouse per group. *: p<0.05. Bar heights and whiskers represent the mean +/- SEM.

The FXS neurons analyzed were matched with FMRP-positive (FMRP+) isogenic control neurons localized in the same brain areas. The maturation stage of FXS and FMRP+ control neurons was assessed by evaluating the fraction of DCX+, DCX+/NeuN+ and NeuN+ in the total of cells staining for DCX or NeuN at 15 days, 1 and 3 months PI. As no expression of DCX was detected in transplanted neurons at 6 months PI, the maturation stage at 6 months PI was assessed using Tuj1, a generic neuronal marker expressed from mid-maturation by neurons and that keeps being expressed in a subpopulation of mature neurons, and NeuN. We determined the percentage of Tuj1+, Tuj1+/ NeuN+ and NeuN+ neurons as compared to the total number of neurons, defined by Tuj1 and/or NeuN labeling.

At 15 days post-injection, the percentage of DCX-positive neurons and the percentage of DCX+NeuN+ neurons were increased and decreased, respectively, in FXS cells as compared to isogenic control suggesting that the maturation of FXS neurons was initially delayed (Figure 3A-C). At 1 month PI, the percentage of neurons labeled with DCX and/or NeuN was similar between FXS and isogenic deletion control, suggesting that there was no difference of maturation between control and FXS neurons for this pair (Figure 3D). In contrast, for the demethylation isogenic pair, less neurons were labeled with NeuN, suggesting that FXS neurons were more immature than control neurons at that stage (Figure 3E). At 3 months PI, however, for both isogenic pairs, the proportion of NeuN+ neurons was increased, indicating more advanced maturation of FXS neurons compared to isogenic control at this later stage (Figure 3F-H). At 6 months PI, virtually all Tuj1 positive cells expressed NeuN, suggesting that FXS and control neurons had reached full maturation at this timepoint. A small but significant increase in the percentage of NeuN+ FXS cells compared to control could still be observed for the demethylation pair, in line with accelerated maturation of FXS neurons (Figure 3I-L).

Together, these observations suggest that the maturation of FXS neurons is, after an initial delay, accelerated compared to isogenic control in the mouse brain. The differences between the two pairs at 1 month and 6 months PI may be explained by the fact the FXS cell line from the demethylation pair (FX52), displays slower neuronal maturation compared to the FXS cell line from the deletion pair (FXS_SW), as indicated by increased percentages of DCX+ neurons and decreased percentages of NeuN+ neurons for the FX52 compared to FXS_SW at all timepoints studied. As a result, the initial delay in maturation persisted at 1 month for FX52 neurons but not for FXS_SW neurons, and accelerated maturation compared to control was still visible at 6 months for FX52 but not for FXS_SW neurons.

### Increased synaptic activity of transplanted FXS neurons

Previous *in vitro* experiments published by our group and other groups showed that FXS neurons in both isogenic pairs were hyperexcitable in vitro ^12, 13^. Immediate early genes (IEG) are expressed during synaptic plasticity and commonly used as markers for synaptic activity. Our single-cell RNA sequencing analysis at 1 month PI showed upregulation of synaptic signaling pathways in FXS neurons, and showed significant upregulation of the IEG ARC, EGR1 and FOS (suppl Figure 10). To confirm that Arc was upregulated in FXS neurons, we used immunofluorescence on transplanted mouse brain slices to assess Arc expression in control and FXS neurons. Neurons were defined by DCX positivity at 1 month PI, as most neurons express DCX at this timepoint and by NeuN staining at 3 months and 6 months PI, as the majority of neurons express this marker at these later timepoints. At 1 month PI, a higher percentage of FXS neurons from the deletion pair was Arc-positive (Arc+) compared to the control (Figure 4A). No significant difference was observed for the demethylation pair, although the percentage of FXS Arc+ neurons was slightly higher than in the isogenic control (Figure 4B). At 3 and 6 months PI, an increased percentage of FXS neurons was labeled with Arc as compared to the controls (Figure 4C-G). Arc is a direct target of FMRP: its translation is repressed by FMRP, and lack of FMRP is expected to increase Arc expression. Therefore, we determined whether Egr1, which is not a direct target of FMRP, was upregulated in FXS neurons, and assessed Egr1 immunolabeling of FXS and control neurons. Similarly to Arc labeling, a higher percentage of FXS neurons was Egr1-positive at all the timepoints studied (Figure 5), further indicating increased activity of FXS neurons compared to control. The IEG c-fos is typically expressed at lower levels in the mouse brain than Arc and Egr1^40–43^, resulting in a low proportion of c-fos positive cells in the brain without stimulation. Out of of 38 to 51 transplanted neurons analyzed, we could not find c-fos-positive control or FXS cells at the timepoints studied.

**Figure 4:**
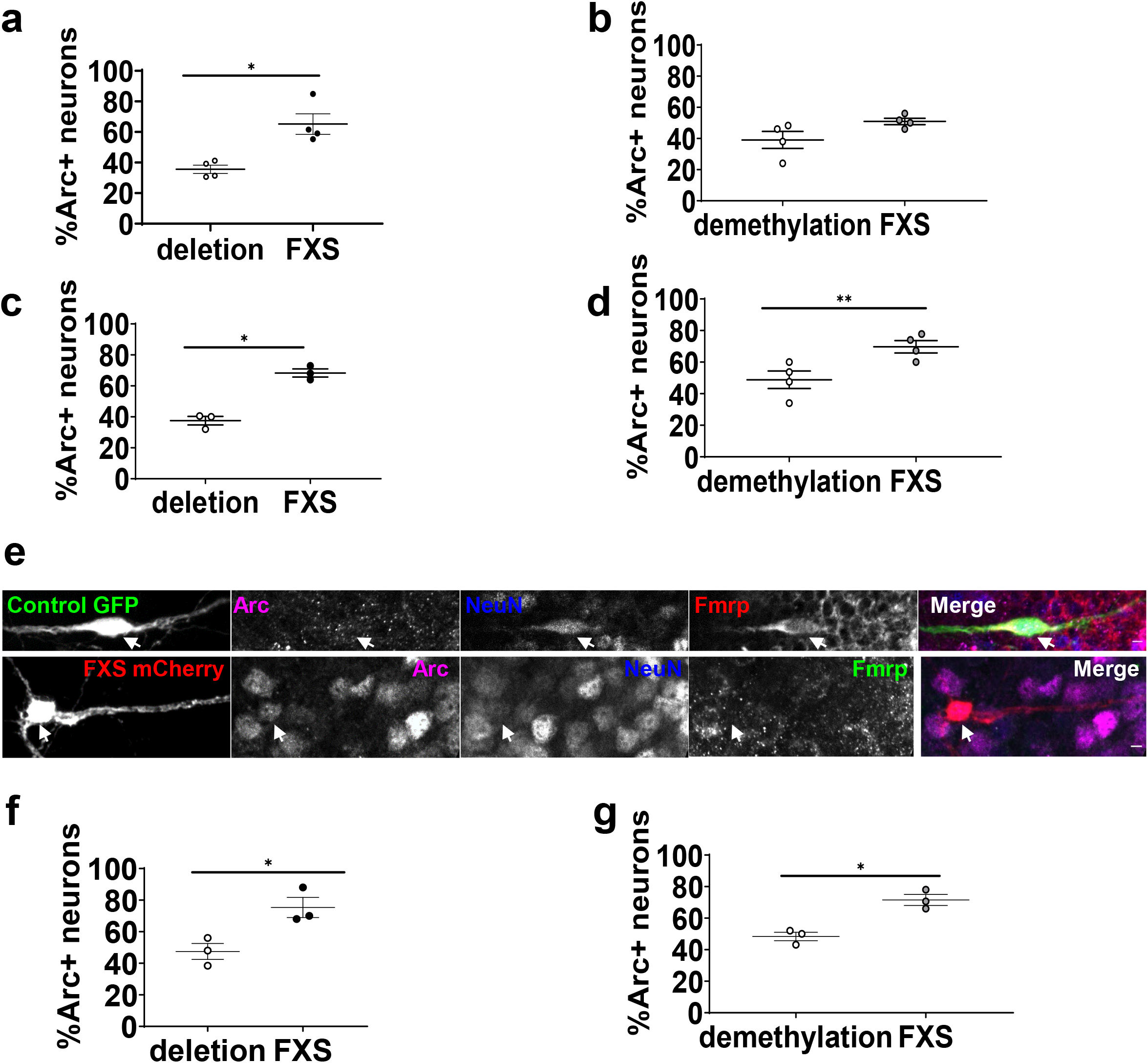
Increased percentages of Arc-positive transplanted FXS neurons from 3 months PI. **a.** Percentage of isogenic control (deletion) and FXS Arc-positive neurons at 1 month PI. Paired t-test; N=4 animals, 36 to 78 neurons analyzed per animal per group. **b.** Percentage of isogenic control (demethylation) and FXS Arc-positive neurons at 1 month PI. Paired t-test; N=4 animals, 50 to 60 neurons analyzed per animal per group. **c.** Percentage of isogenic control (deletion) and FXS Arc-positive neurons at 3 months PI. Paired t-test; N=3 animals, 37 to 50 neurons analyzed per animal per group. **d.** Percentage of isogenic control (demethylation) and FXS Arc-positive neurons at 3 months PI. Paired t-test; N=4 animals, 50 to 61 neurons analyzed per animal per group. **e.** Confocal plane showing an isogenic control (demethylation) Fmrp-positive Arc-negative neuron and an FXS Fmrp-negative Arc-positive neuron at 3 months PI. White arrows indicate neuronal cell bodies. Scale bars represent 5 µm. **f.** Percentage of isogenic control (deletion) and FXS Arc-positive neurons at 6 months PI. Paired t-test; N=3 animals, 50 to 52 neurons analyzed per animal per group. **g.** Percentage of isogenic control (demethylation) and FXS Arc-positive neurons at 6 months PI. Paired t-test; N=3 animals, 50 to 51 neurons analyzed per animal per group. Neurons were defined as doublecortin-positive cells at 1 month post-injection (PI) and as NeuN-positive cells at 3 and 6 months PI. *: p<0.05; **: p<0.01. Thin horizontal bars in A-D, F, G indicate the mean.

**Figure 5:**
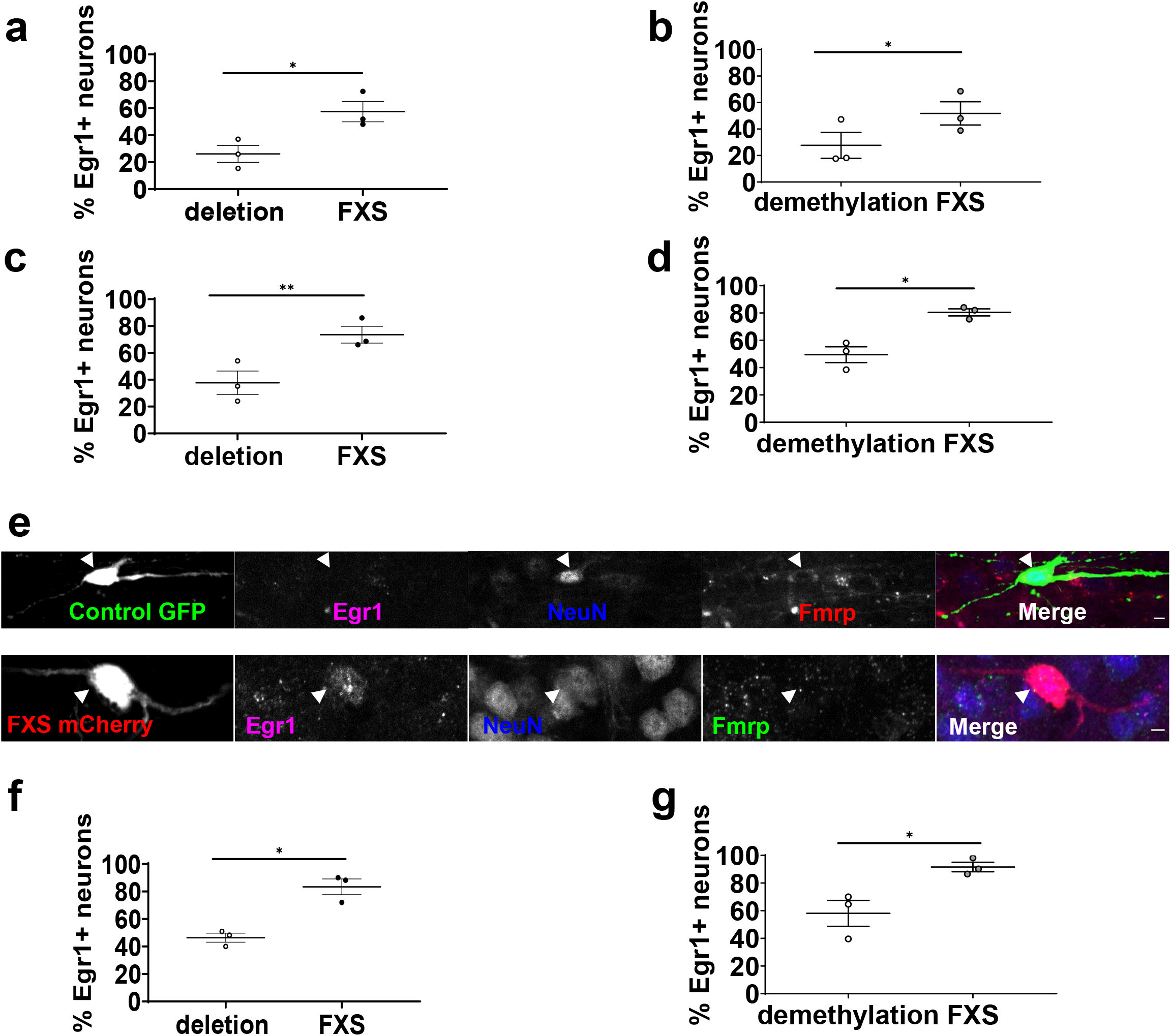
Increased proportions of Egr1-positive transplanted FXS neurons at all the timepoints studied. **a.** Percentage of isogenic control (deletion) and FXS Egr1-positive neurons at 1 month PI. Paired t-test; N=3 animals, 51 to 54 neurons analyzed per animal per group. **b.** Percentage of isogenic control (demethylation) and FXS Egr1-positive neurons at 1 month PI. Paired t-test; N=3 animals, 52 to 55 neurons analyzed per animal per group. **c.** Percentage of isogenic control (deletion) and FXS Egr1-positive neurons at 3 months PI. Paired t-test; N=3 animals, 50 to 54 neurons analyzed per animal per group. **d.** Percentage of isogenic control (demethylation) and FXS Egr1-positive neurons at 3 months PI. Paired t-test; N=3 animals, 50 to 53 neurons analyzed per animal per group. **e.** Confocal plane showing an isogenic control (demethylation) Fmrp-positive Egr1-negative neuron and an FXS Fmrp-negative Egr1-positive neuron at 3 months PI. White arrows indicate neuronal cell bodies. Scale bars represent 5 µm. **f.** Percentage of isogenic control (deletion) and FXS Egr1-positive neurons at 6 months PI. Paired t-test; N=3 animals, 50 to 56 neurons analyzed per animal per group. **g.** Percentage of isogenic control (demethylation) and FXS Egr1-positive neurons at 6 months PI. Paired t-test; N=3 animals, 50 to 53 neurons analyzed per animal per group. Neurons were defined as doublecortin-positive cells at 1 month post-injection (PI) and as NeuN-positive cells at 3 and 6 months PI. *: p<0.05; **: p<0.01. Data is presented as mean±SEM.

To further investigate lasting changes in arborization complexity, excitatory synaptogenesis and synaptic activity in FXS neurons, we assessed the dendritic protrusion density and morphology of mature FXS and isogenic control neurons in the striatum at 6-7 months PI. As the striatum is the brain region with the most transplanted human neurons still present at 6 months PI, and as mature medium spiny neurons display complex morphology and high dendritic spine density, we focused our analysis on the medium spiny neurons of the striatum. We used DARPP32 as a marker for mature medium spiny neurons (MSNs) (Suppl. Figure 11A). Control and FXS MSNs exhibited complex morphology at 6-7 months PI (Suppl. Figure 11B, C). No difference in neuronal arborization complexity was detected between FXS and control MSNs (Suppl. Figure 11B-F). Dendritic spines generally correspond to excitatory synapses^44, 45^. It is commonly accepted that increased spine head size correlates with increased synaptic strength, and dendritic spines have been found to become wider after long-term potentiation ^46^. We measured dendritic protrusion density and dendritic protrusion head diameter in control and FXS neurons, using automated tracing and analysis with the Imaris software. Dendritic protrusion end diameter was significantly increased in FXS MSNs for both pairs (Figure 6A-D). We classified dendritic protrusions into three categories in function of the size of their head: head size inferior or equal to 0.3 µm, head size strictly comprised between 0.3 and 0.6 µm, and head size ≥ 0.6 µm. For both pairs, FXS neurons had more large protrusions (head size superior or equal to 0.6 µm) and less thin protrusions (diameter between 0.3 and 0.6 µm) than control neurons (Figure 6A, B, E, F). No significant change in dendritic protrusion density could be detected between control and FXS neurons (Figure 6G,H). This observation suggests that, although excitatory synaptogenesis in FXS MSNs is unchanged, excitatory synaptic strength is increased, which is in line with the increased excitatory synaptic activity in FXS neurons we observed. Together, these results further indicate increased synaptic activity, but not synaptogenesis, in FXS neurons compared to control.

**Figure 6:**
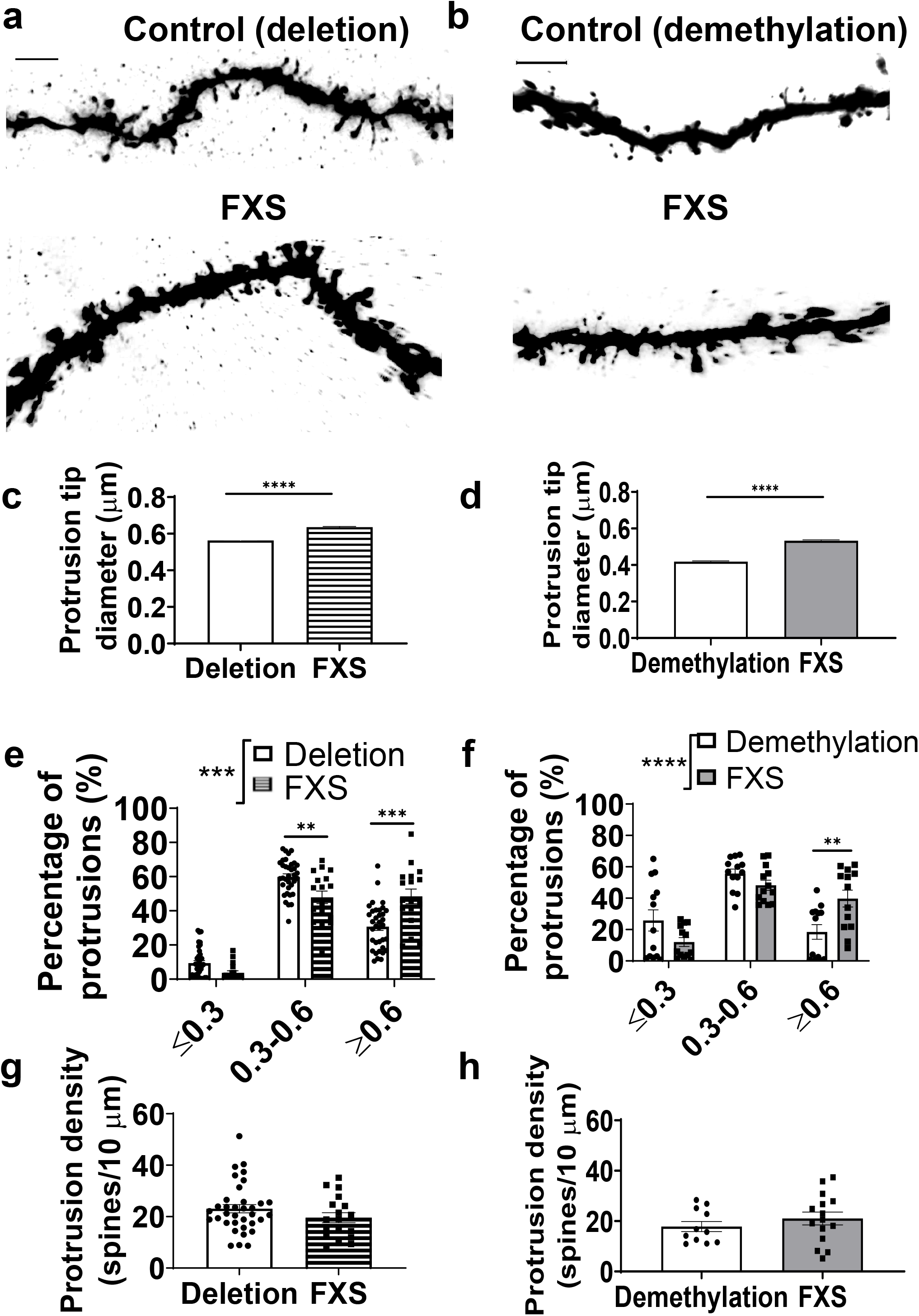
Wider dendritic protrusions in transplanted FXS striatal medium spiny neurons (MSNs) at 6-7 months post-injection. **a**. Representative 3D reconstructions of dendritic segments of control (deletion) (**upper panel**) and FXS (**lower panel**) MSNs. **b.** Representative 3D reconstructions of dendritic segments of control (demethylation) (**upper panel**) and FXS (**lower panel**) MSNs. Average tip diameter of dendritic protrusions of control and FXS MSNs from the deletion (**c**) and demethylation (**d**) pairs. Unpaired Mann-Whitney test; N=1945 to 4509 protrusions from 13 to 35 MSNs per group. Percentage of protrusions with tip diameter inferior or equal to 0.3 µm (≤0.3), strictly comprised between 0.3 and 0.6 µm (0.3-0.6), or superior or equal to 0.6 µm (≥0.6) in control and FXS MSNs from the deletion (**e**) and demethylation (**f**) pairs. Mixed-effects model followed by Sidak’s correction for multiple comparisons; N=13 to 35 MSNs per group. Average protrusion density of FXS and control MSNs from the deletion (**g**) and demethylation (**h**) pairs. N=11 to 35 MSNs per group. *: p<0.05; **: p<0.01; ***: p<0.001; ****: p<0.0001. Bar heights and whiskers represent the mean +/- SEM. Scale bars represent 50 µm.

## Discussion

In this study, we transplanted FXS and isogenic control hNPCs in the mouse brain, allowing for neuronal development in the *in vivo* context of the mouse brain. The transplanted hNPCs were sorted for high PSA-NCAM expression but not directed towards a specific neuronal subtype, and gave rise to a high number of neurons at 1 and 3 months PI. Neurons were fully mature at 6 months PI as assessed by DCX and NeuN expression, and some neurons differentiated in MSNs in the striatum, displaying complex morphology characteristic of striatal MSNs, a high dendritic spine density, and expressing DARPP32, a marker specific to striatal MSNs.

We found an initial delay followed by an accelerated maturation of FXS neurons as compared to control. *In vitro* studies have found impaired maturation of FXS neurons^7, 8, 10, 11^. This discrepancy may be explained by the fact that neurons transplanted in the brain of mice reach a higher differentiation stage than neurons grown *in vitro*. Indeed, transplanted neurons reached a high arborization complexity and some formed dendritic spines, in constrast to the neurons grown in the 2D cultures used in these studies, which have less complex arborization and do not form dendritic spines. Therefore, the differentiation stage of neurons in 2D cultures might not be sufficient to reveal the subsequent acceleration in the maturation observed in FXS neurons growing in the mouse brain.

FXS MSNs displayed an increased dendritic protrusion head diameter as compared to isogenic controls in line with the upregulation of synaptic activity pathways as assessed by single-cell RNA sequencing and Arc upregulation. This finding is not in agreement with previous studies in a FXS mouse model that showed dendritic protrusion elongation and no change in dendritic protrusion head diameter in mature striatal MSNs ^47^. Similarly, dendritic spine analysis in FXS patients showed increased spine density and increase in the proportion of spines with an immature morphology, i.e. with a longer neck and smaller head, in neocortical pyramidal cells ^6^. This difference may be explained by a non-cell autonomous effect on the dendritic spine phenotype, possibly caused by FXS astrocytes. In contrast, the transplanted FXS and control neurons in our study were exposed to the same neurodevelopmental niche and, although FXS glial cells were present in the mouse brain and proliferated between 1 and 6 months PI, they were sparse and FXS neurons were not in direct contact with them. Therefore, the effects we observed were most likely cell-autonomous. Consistent with this hypothesis, adult astrocyte-specific *FMR1* KO mice displayed increased spine density and a higher proportion of dendritic spines in the motor cortex, indicating that FXS astrocytes are necessary and sufficient for the dendritic spine phenotype of neurons in FXS ^48^.

Our data on Arc, Egr1 and c-fos expression, along with the percentages of Arc+ and Egr1+ neurons and dendritic protrusion diameter, indicate that FXS neurons display hyperactive synapses compared to controls. This observation is consistent with previous work indicating that the excitation/inhibition balance is altered in FXS patients, in agreement with previous studies from our laboratory and others ^12–14^ that revealed hyperexcitability in FXS neurons grown in 2D cultures. Interestingly, neurons develop to form hyperexcitable networks in other intellectual disability and autism spectrum disorders, such as Kleefstra syndrome, Angelman syndrome and idiopathic ASD ^49–52^.

Two genes of interest, OTX1 and TBX1 were strongly downregulated or upregulated, respectively, in FXS transplanted neurons at 1 month PI, as assessed by scRNA seq (Tables S1 and S2). OTX1 is associated with ASD-associated behavior and epilepsy, two symptoms of FXS. Additionally, OTX1 knock-down was previously shown to increase the astrocyte/neuron ratio in the developing cortex, and to maintain neural progenitors in the transient amplifying phase ^53^. OTX1 expression is also required for the refinement of axonal projections of cortical neurons to subcortical targets ^54^. The downregulation of OTX1 we observed in FXS neurons with phenotypes previously described in FXS models: increased neural progenitor proliferation and aberrant axogenesis ^55–58^. TBX1 is also associated with ASD and intellectual disability. In particular, TBX1 mutations leading to a gain of function were linked to intellectual disability ^59, 60^. This suggests that OTX1 and TBX1 may be interesting targets for rescuing the cell-autonomous phenotypes of FXS neurons.

A major challenge in drug development is assessing the ability of the molecules to pass the blood-brain barrier. Despite its limitations due to the fact that FXS patient-derived transplanted neurons grow among wild-type mouse neural cells, our human-mouse chimeric model provides us with a readout to test potential therapeutic molecules acting on the phenotypes of neurons while accounting for their potential modifications in the organism and their ability to cross the blood-brain barrier, in contrast with 2D or 3D culture models. Therefore, it could be used in addition to 2D and 3D culture models for drug testing. It is of note that our NPC differentiation and transplantation protocol needs to be refined: the number of transplanted neurons decreased between 3 and 6 months PI and transplanted neurons were sparse at 6 months PI. Moreover, glial cells proliferated over time, becoming more numerous between 1 and 6 months PI. These factors made the analysis at later time points challenging. Additionally, our approach does not allow us to control the fate of the transplanted cells, making the study of different neuronal subtypes difficult. To solve this issue, one could consider injecting hNPCs directed towards a striatal, cortical or hippocampal fate, or, as done in previous studies, cortical or striatal neurons ^61–63^. A potential limitation of the neuronal injection approach is the lower contribution of the cells to the mouse brain at early timepoints, as neurons are less resistant to stress than neural precursor cells and do not proliferate after injection.

## Acknowledgements

We thank Patti Wisniewski, Patrick Autissier, and Hanna Aharonov from the FACS facility of Whitehead Institute for Biomedical Research for FACS, Jennifer Love from the Genomics core of Whitehead Institute for single-cell sequencing service. We thank George Bell for helpful discussions about statistics. We thank Wendy Salmon from the Keck Microscopy Facility of Whitehead Institute for her useful suggestions on confocal microscopy acquisitions. We thank Steve Warren in Emory University for providing FXS_SW isogenic iPSC lines. We thank members of the Jaenisch lab and Fulcrum Therapeutics for discussion and suggestions on the manuscript. We thank Li-Huei Tsai and her group for providing the dissociator used for the extraction of transplanted cells. This data was presented during the International Society for Stem Cell Research (ISSCR) virtual meeting 2020.

## Author Contributions

M.K., H.W., A.C. and R.J. conceived the idea for this study. M.K. and H.W. designed the experiments. M.K. interpreted the data. B.Y. and T.W.W. analyzed the single-cell sequencing data. M.K., H.W., D.F., C.N., J.S. performed the experiments. M.K., C.M.G., K.R.A, R.R., J.S. and B.J. analyzed the data. S.L. provided FXS2 and isogenic control iPSCs. S.W. provided the FXS_SW and isogenic control iPSCs. O.W. oversaw the first part of the study. M.K. and R.J. wrote the manuscript with input from all the other authors.

