## Supplemental material and table legends for "Fragile X syndrome patient-derived neurons developing in the mouse brain show *FMR1* -dependent phenotypes"

### Supplementary figure and table legends

**Suppl. Figure 1**: Discrepancy between $\log_2$ fold change ($\log_2$FC) with p-value for scarcely expressed genes in neurons (**a**) and NPC (**b**). Genes expressed in less than 1% of cells are highlighted in red. The blue lines marked the two fold differences or adjusted p-value equaled to 0.05.

**Suppl. Figure 2**: **Isogenic pairs used in this study.** Two different male FXS cell lines were used. In FXS_SW, CGG repeats were deleted using CRISPR editing, and in FXS2, CGG repeats were demethylated using dCas9-Tet1/single guide RNA. Both approaches led to the reactivation of the Fmr1 promoter.

**Suppl. Figure 3: hNPCs derived from isogenic control and FXS iPSC express neural progenitor markers.** Confocal maximum intensity projections of immunostainings against Nestin (green), Sox2 (red) and and Pax6 (magenta) on cultured hNPCs from the different isogenic pairs. Scale bars represent 20 µm.

**Suppl. Figure 4: No doublecortin/Olig2 co-staining was found in the transplanted cells. a.** Confocal maximum intensity projection showing Olig2-positive doublecortin (DC)-negative cells. **b.** Confocal maximum intensity projection showing doublecortin-positive Olig2-negative cells. White arrows indicate neuronal cell bodies. Scale bars represent 20 µm.

**Suppl. Figure 5: Fluorescent cells in the chimeric brains are all of human origin.** Representative confocal maximum intensity projections of the cortical region of a mouse brain at 1 month post-injection engrafted with tdTomato-labeled C1_2_SW and GFP-labeled FXS_SW isogenic NPCs at P1. All the engrafted cells were stained for human nuclei antigen (HuNu) and GFP+ and tdT+ cells were mutually exclusive. Stained in green, GFP; red, tdT; magenta, huNu; blue, DAPI.

**Suppl. Figure 6: Isogenic control cells transplanted in the mouse brain express Fmrp. a.** Confocal maximum intensity projections of cell bodies of control (demethylation) GFP Fmrp-positive transplanted NeuN-positive neurons (green, upper panel) and FXS tdT transplanted Fmrp-negative NeuN-positive neurons (red, lower panel) at 3 months PI. White arrows indicate neuronal cell bodies. Scale bars represent 5 µm. **b.** Percentage of isogenic control (deletion) neurons expressing Fmrp as assessed by immunofluorescence against Fmrp at 1, 3 and 6 months PI. Neurons were defined as doublecortin-positive cells at 1 month PI and NeuN-positive cells at 3 and 6 months PI. One-way ANOVA; N=3 to 4 animals, 25 to 59 neurons analyzed per animal per group. **c.** Percentage of isogenic control (demethylation) neurons reexpressing Fmrp as assessed by immunofluorescence against Fmrp at 1, 3 and 6 months PI. Neurons were defined as doublecortin-positive cells at 1 month PI and NeuN-positive cells at 3 and 6 months PI. One-way ANOVA; N=3 to 4 animals, 50 neurons analyzed per animal per group. **d.** *FMR1* expression by RT-qPCR on total mRNA of isogenic control (deletion) and FXS cells extracted from the mouse brain at 1 month PI. Unpaired t-test, N= 3 technical replicates. **e.** DNA methylation level of 10 CpGs in the promoter of FMR1 measured by Pyro-sequencing of transplanted cells extracted from the mouse brain at one-month PI. **f.** Average DNA methylation levels of the 10 CpGs in the promoter of FMR1 presented in **g.** Bar heights and whiskers represent the mean +/- SEM. ***: $p<0.001$

**Suppl. figure 7: Characterization of UMAP clusters. a.** DCX expression levels in UMAP clusters. More mature neurons still express DCX. **b.** Cell cycle markers in the different UMAP clusters. Imm: immature; M. m.: more mature. **c.** Zoomed-in part of the heatmap for transplanted cells, featuring the 15 cells from the cluster More mature neurons 2. **d.** Pseudotime analysis of the UMAP clusters with Slingshot without specifying a starting cluster. Imm: immature. **e.** Two lineages are predicted by Slingshot with NPC 1 as the starting point (green dot). $1^{st}$ lineage: NPC -> Immature neurons to more mature neurons. $2^{nd}$ lineage: NPC to glial cells. Cells are showed in the principle component 1 (PC 1) and the principle component 2 (PC 2).

**Suppl. figure 8: FMR1 is down-regulated in FXS cells**. After randomly selecting 1000 cells from each sample, normalized counts were plotted only for cells expressing FMR1. Points with the same normalized expression were shifted left or right to avoid overlaps in the visualization. FMR1 expression was significantly lower in FXS samples versus isogenic controls with FDR < 9.4e-27 using edgeR on aggregated counts.

**Suppl. figure 9: Single cell RNA seq analysis suggests increased maturation of FXS neurons at 1 month post-injection. a.** Inverted beta-binomial test for paired count data. The cell counts were normalized so that each sample had the same total counts. The distribution of FXS cells is shifted towards more mature clusters compared to isogenic control. A positive value in fold change (FC) means up-regulation in FXS, and a negative value down-regulation in FXS. Gene Set Enrichment Analysis (GSEA) reveals no significant difference between control and FXS NPCs **(b)** and neurons **(c**) in the expression of hallmark apoptosis genes sets from MSigDB. FDR q-value NPC: 0.05. FDR q-value neurons: 0.9. d. Violin plots shows that the percentages of mitochondrial genes are similar in control and FXS cells.

**Suppl. figure 10: Single cell RNA sequencing shows upregulation of ARC, FOS and EGR1 in FXS neurons**

**Suppl. figure 11: Control and FXS striatal medium spiny neurons (MSNs) do not display significant changes in neuronal arbor complexity at 6-7 months post-transplantation. a.** Single confocal planes of FXS (tdT) and isogenic control (GFP) neurons stained with DARPP32. **b.** Representative confocal maximum intensity projections and corresponding tracings of transplanted control (left panel) and FXS (right panel) DARPP32-positive neurons from the deletion pair. **c.** Representative 3D reconstructions and corresponding tracings of transplanted control (left panel) and FXS (right panel) DARPP32-positive neurons from the demethylation pair. **d.** Scheme illustrating bifurcations (nodes), terminals, cables (branches) and stems of a neuron. **e.** Number of bifurcations, cables, stems and terminals of FXS and isogenic control (deletion) DARPP32-positive control neurons. Mixed-effects analysis; N=14 to 27 neurons per group. **f.** Number of bifurcations, cables, stems and terminals of FXS and isogenic control (demethylation) DARPP32-positive control neurons. Mixed-effects analysis; N=11 to 14 neurons per group. Scale bars represent 20 μm.

**Table S1: Most strongly downregulated genes in FXS neurons as assessed by single cell RNA seq analysis.** The threshold was set at $\log_2 FC < -2$

**Table S2: Most strongly upregulated genes in FXS neurons as assessed by single cell RNA seq analysis.** The threshold was set as $\log_2 FC > 2$.

**Table S3: FXS patient-derived iPSC lines and isogenic control cell lines used in this study**

**Table S4: Antibodies used in this study**
