## Supplemental tables for "Fragile X syndrome patient-derived neurons developing in the mouse brain show *FMR1* -dependent phenotypes"

### Supplementary tables

### Table S1

| Gene symbol | Gene name | Function | $Log_2$(FC) |
|---|---|---|---|
| **FMR1** | FMRP translational regulator 1 | Translational regulator | -7.0 |
| **CA9** | Carbonic anhydrase 9 | Carbonic anhydrase | -6.3 |
| **HHEX** | Hematopoietically expressed homeobox | Transcription factor | -6.2 |
| **ZNF736** | Zinc finger protein 736 | Involved in transcriptional regulation? | -3.0 |
| **HES5** | Hes family bHLH transcription factor 5 | Transcription factor | -2.5 |
| **OTX1** | Orthodenticle homeobox 1 | Transcription factor | -2.1 |
| **CDCA2** | Cell division cycle associated 2 | Regulation of DNA damage response | -2.1 |

### Table S2

| Gene symbol | Gene name | Function | $Log_2$(FC) |
|---|---|---|---|
| **HOXC10** | Homeobox C10 | Transcription factor | 8.9 |
| **TAF9B** | TATA-box binding protein associated factor 9b | Transcription factor subunit | 6.8 |
| **NKX2-2** | NK2 homeobox 2 | Transcription factor | 6.4 |
| **TBX1** | T-box transcription factor 1 | Transcription factor | 5.8 |
| **HOXA10** | Homeobox A10 | Transcription factor | 4.3 |
| **CCDC125** | Coiled-coil domain containing 125 | Regulation of cell migration? | 4.1 |
| **SPINK5** | Serine peptidase inhibitor Kazal type 5 | Serine protease inhibitor | 4.0 |
| **C9orf64** | Chromosome 9 open reading frame 64 | Unknown | 3.8 |
| **HMX1** | H6 family homeobox 1 | Transcription factor | 3.5 |
| **TRIM61** | Tripartite motif containing 61 | unknown | 3.2 |
| **SP140L** | SP140 nuclear body protein like | unknown | 3.1 |
| **HOXD3** | Homeobox D3 | Transcription factor | 3.1 |

### Table S3

| Induced pluripotent stem cell lines | Reference |
|---|---|
| **FXS_SW (from Steven Warren's group)** | Xie et al., 2016 |
| **C1_2_SW (from Steven Warren's group)** | Xie et al., 2016 |
| **FXS2** | Park et al., 2015 |
| **FXS2 dCT** | Liu et al., 2018 |
| **FXS2 dCdT** | Liu et al., 2018 |

### Table S4

| Reagent | Source | Identifier |
|---|---|---|
| **Primary antibodies** | | |
| Guinea pig anti-Arc | Synaptic Systems | 156 004 |
| Guinea pig anti-doublecortin | MilliporeSigma | AB2253MI |
| Rabbit anti-doublecortin | Cell Signaling Technology | 4604 |
| Rat anti-DARPP32 | Lifespan Biosciences | LS-C36138 |
| Rat anti-Egr1 | Lifespan biosciences | LS-C36221 |
| Rabbit anti-FMRP | Cell Signaling Technology | 4317S |
| Rabbit anti-FMRP | Cell Signaling Technology | 7104 |
| Chicken anti-GFP | AVES Labs | GFP-1020 |
| Mouse anti-human nuclei | MilliporeSigma | MAB1281 |
| Guinea pig anti-NeuN | Life Technologies | ABN90MI |
| Mouse anti-NeuN | MilliporeSigma | MAB377 |
| Goat anti-tdTomato | SICGEN | AB8181-200 |
| Mouse anti-Tuj1 | Biolegend | 801201 |
| **Secondary antibodies** | | |
| Donkey Alexa 488 fluor anti-chicken | Jackson ImmunoResearch | 703-545-155 |
| Donkey Alexa 555 fluor anti-goat | Life Technologies | A21432 |
| Donkey IRDye 680LT anti-guinea pig | LI-COR Biosciences | 925-68030 |
| Donkey Alexa fluor 405 anti-mouse | Abcam | ab175659 |
| Donkey Alexa fluor 647 anti-mouse | Life Technologies | A31571 |
| Donkey Alexa fluor 405 anti-rabbit | Fisher Scientific | NC0192764 |
| Donkey Alexa fluor 488 anti-rabbit | Life Technologies | A21206 |
| Donkey Alexa fluor 594 anti-rabbit | Life Technologies | A21207 |
| Donkey Alexa fluor 647 anti-rabbit | Life Technologies | A31573 |
| Donkey Alexa fluor 647 anti-rat | Jackson ImmunoResearch | 712-605-153 |
