## Supplemental material for "Fragile X syndrome patient-derived neurons developing in the mouse brain show *FMR1* -dependent phenotypes"

### Supplementary material

### Suppl. Figure 1

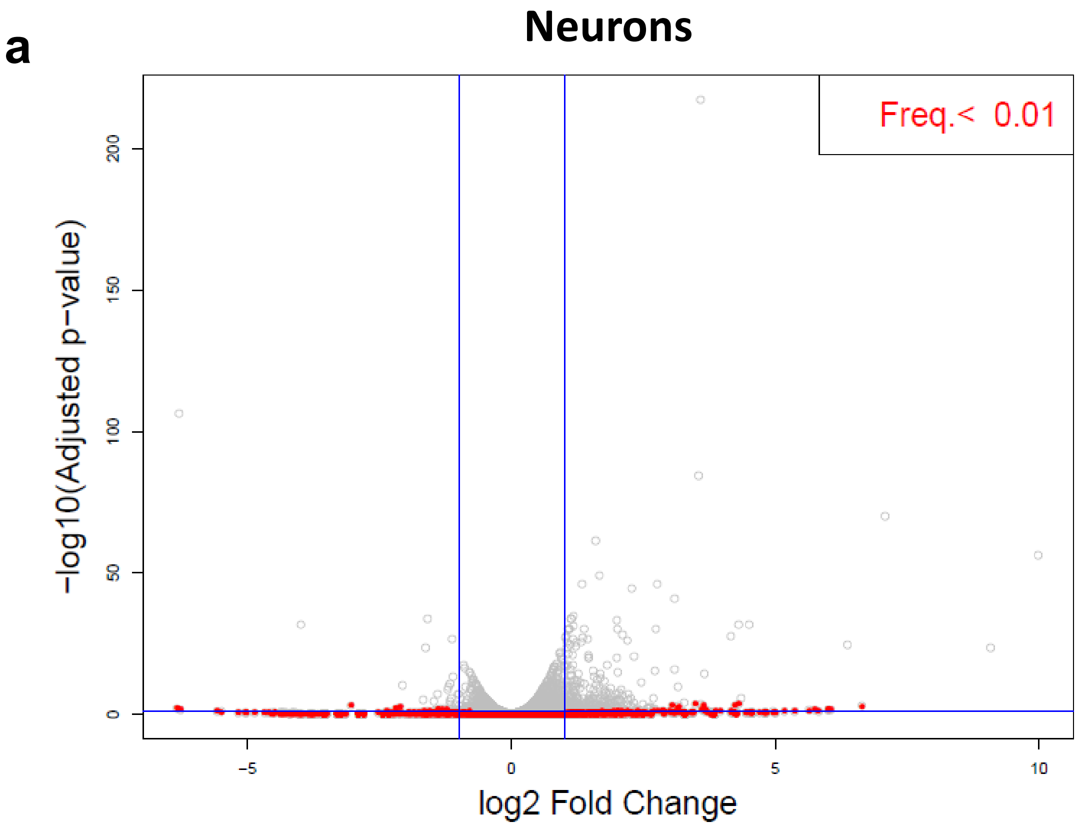


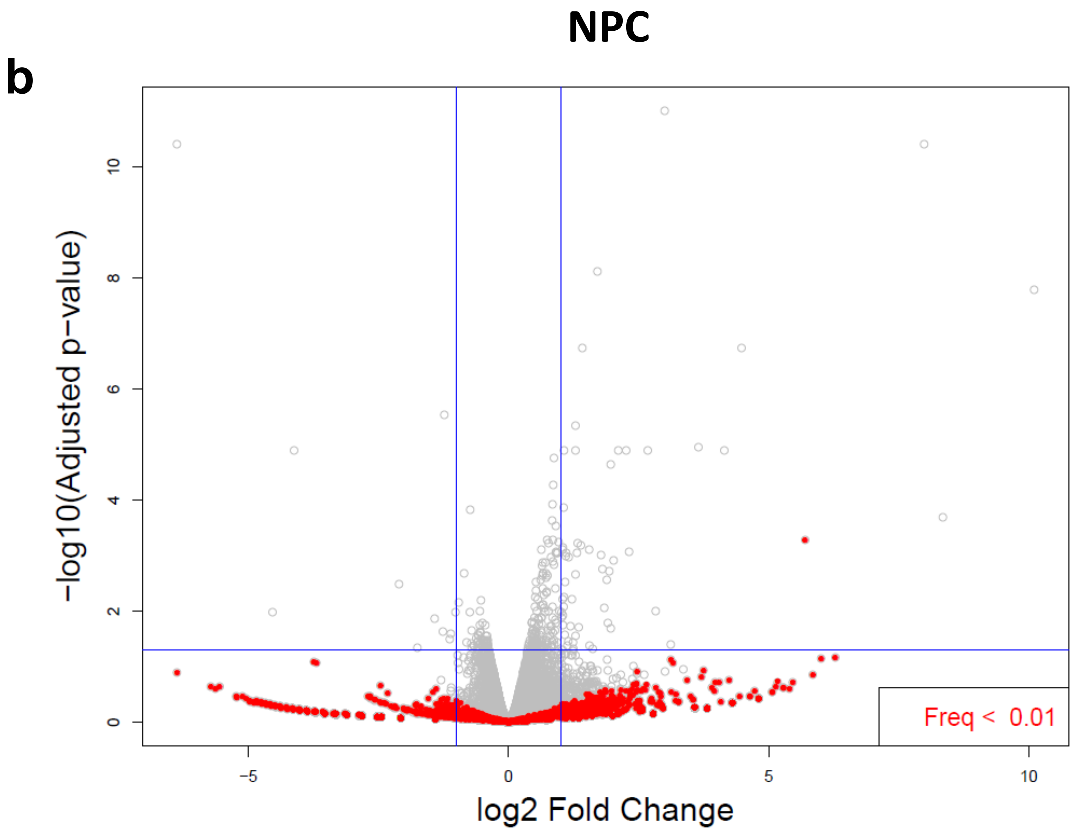

### Suppl. Figure 2

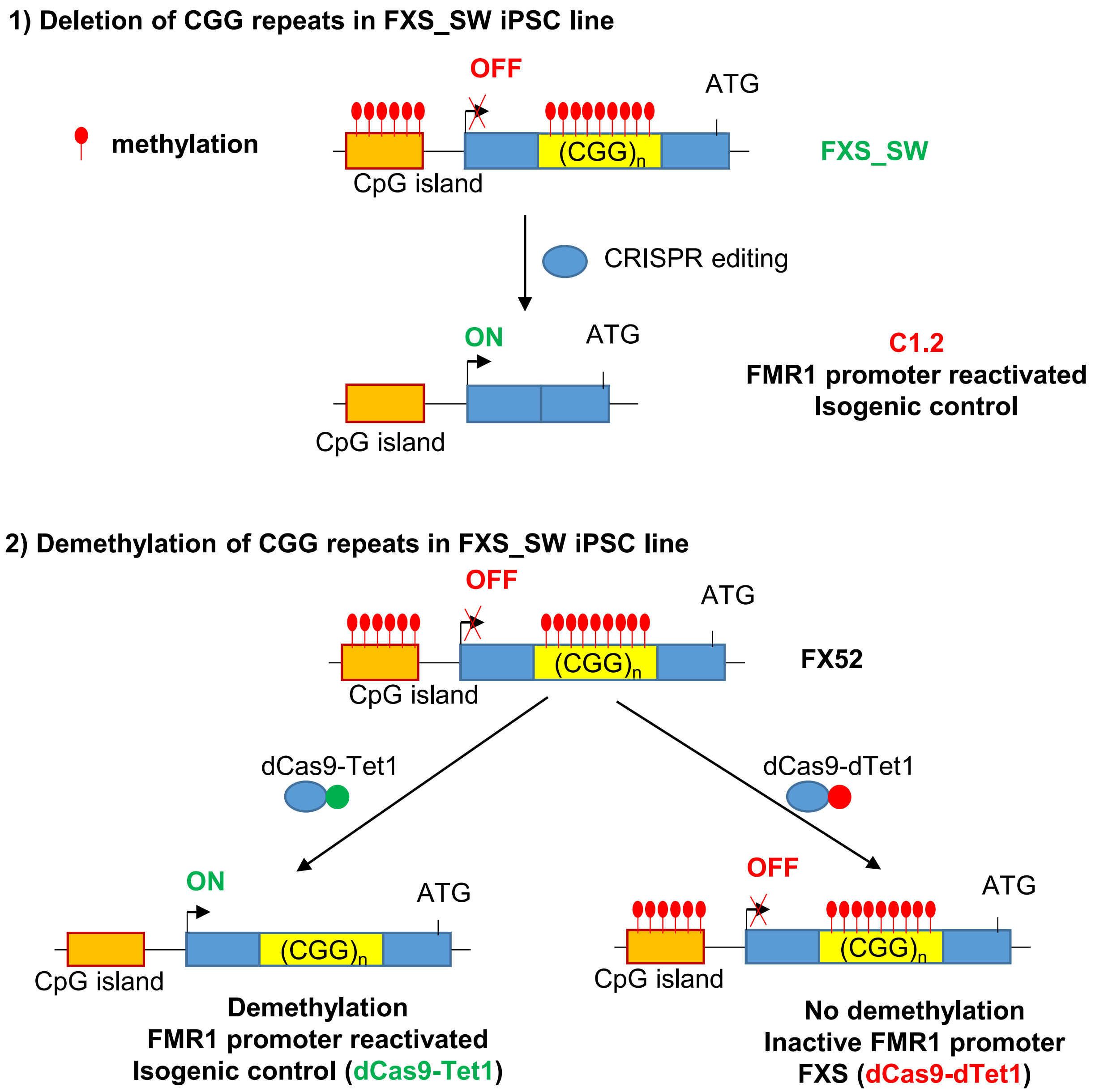

### Suppl. Figure 3

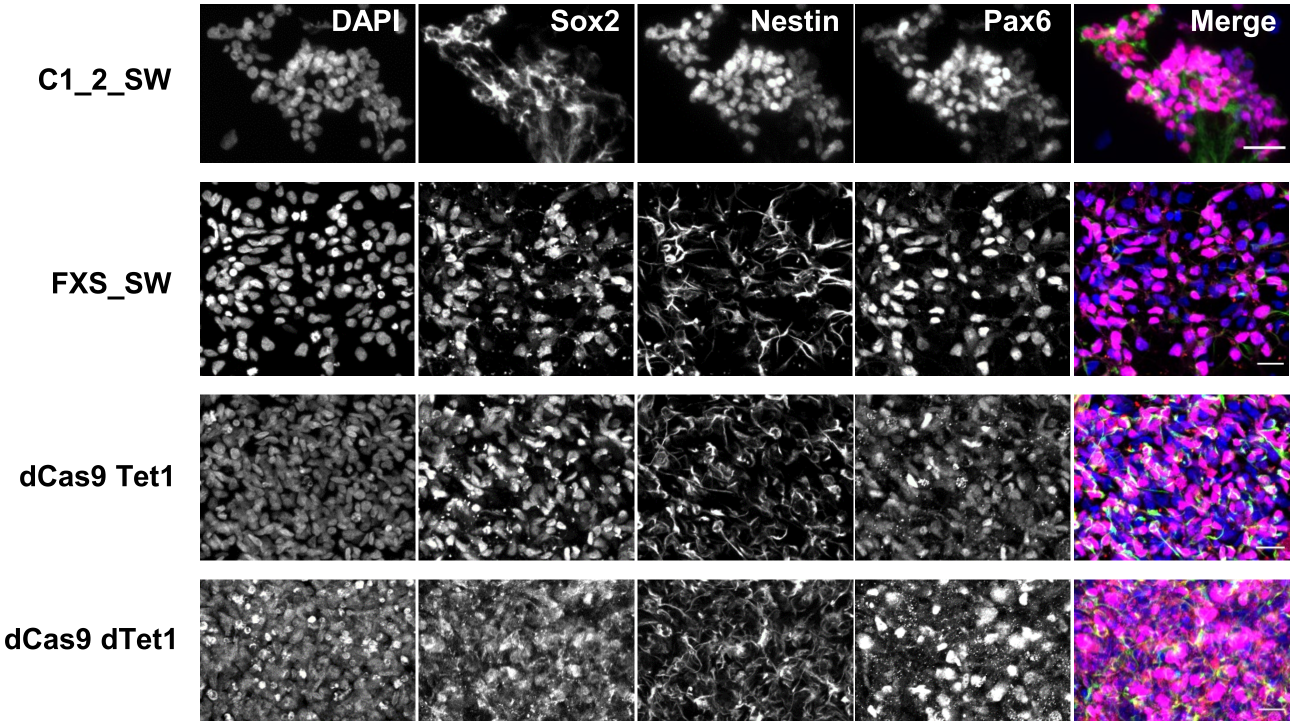

### Suppl. Figure 4

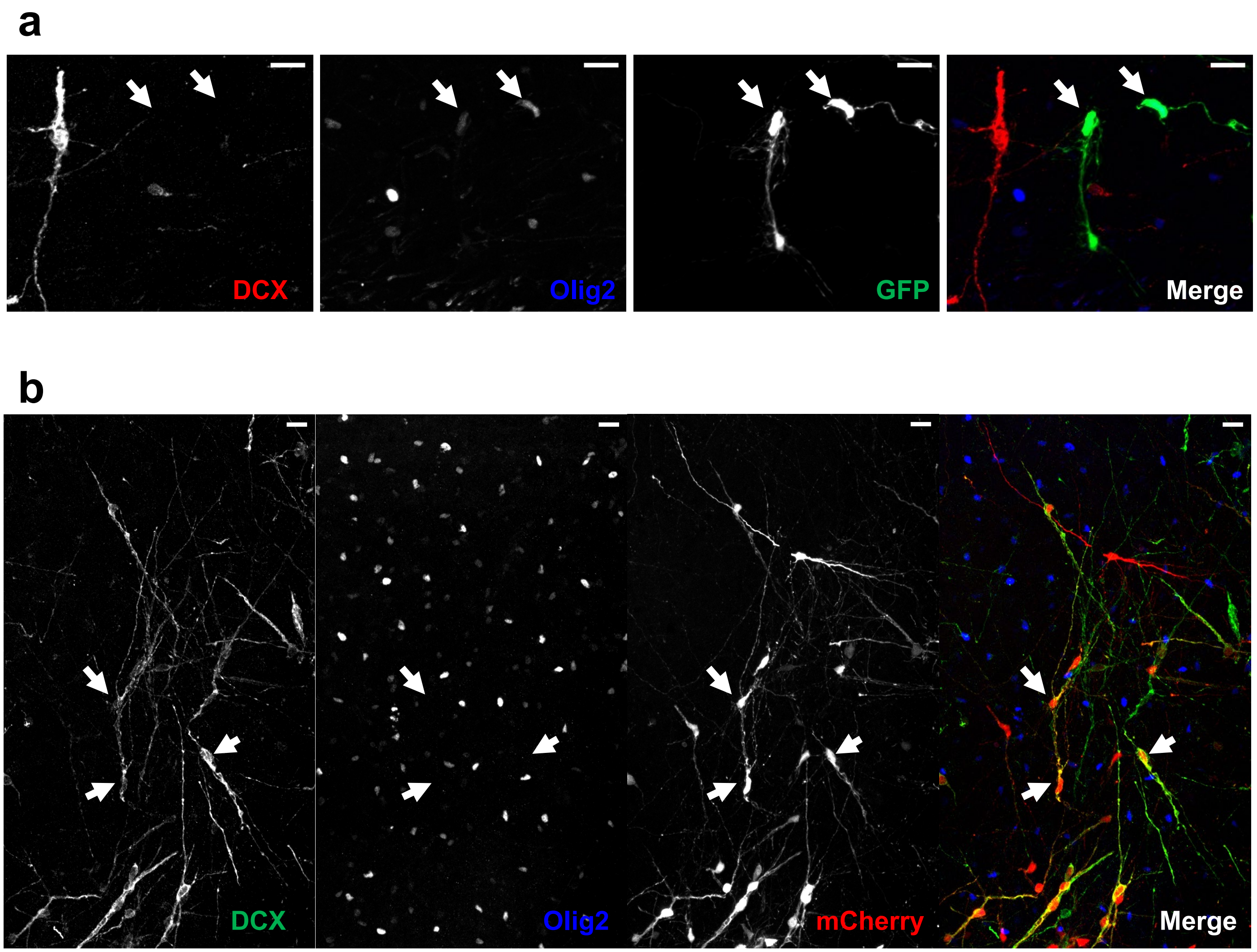

### Suppl. Figure 5

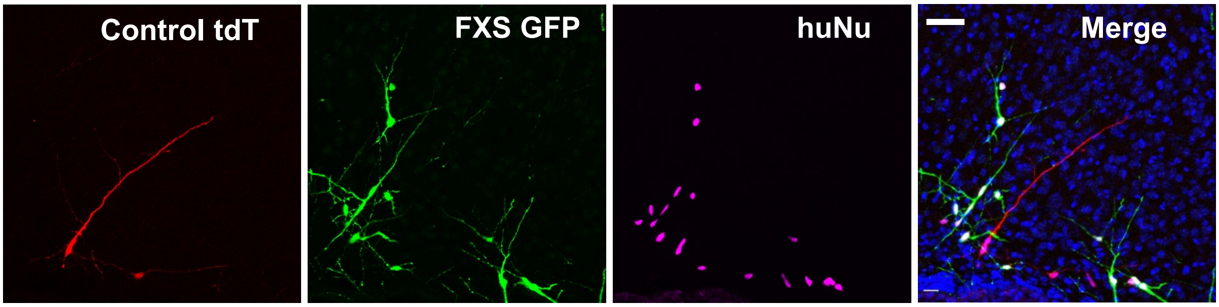

### Suppl. Figure 6

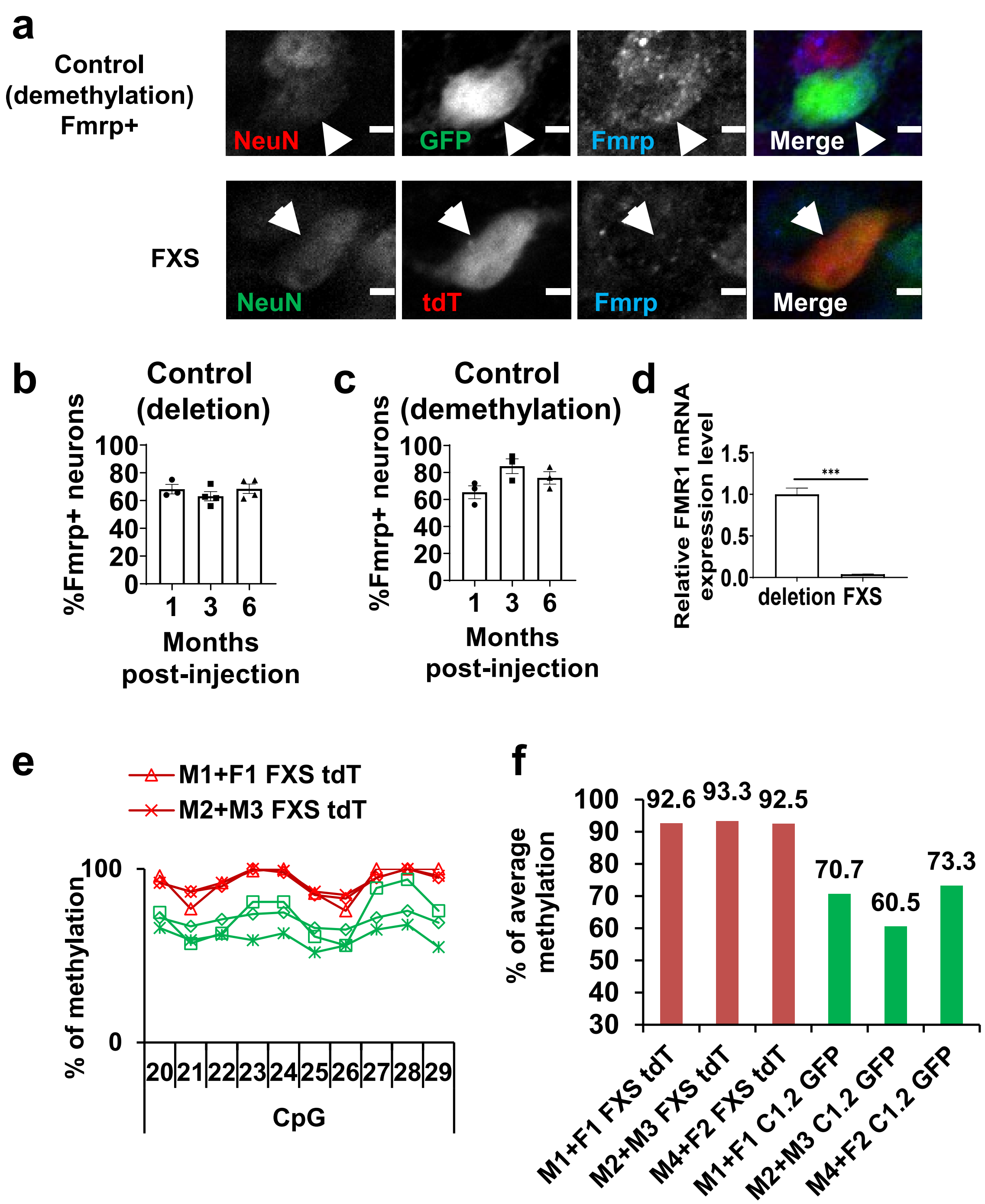

### Suppl. Figure 7

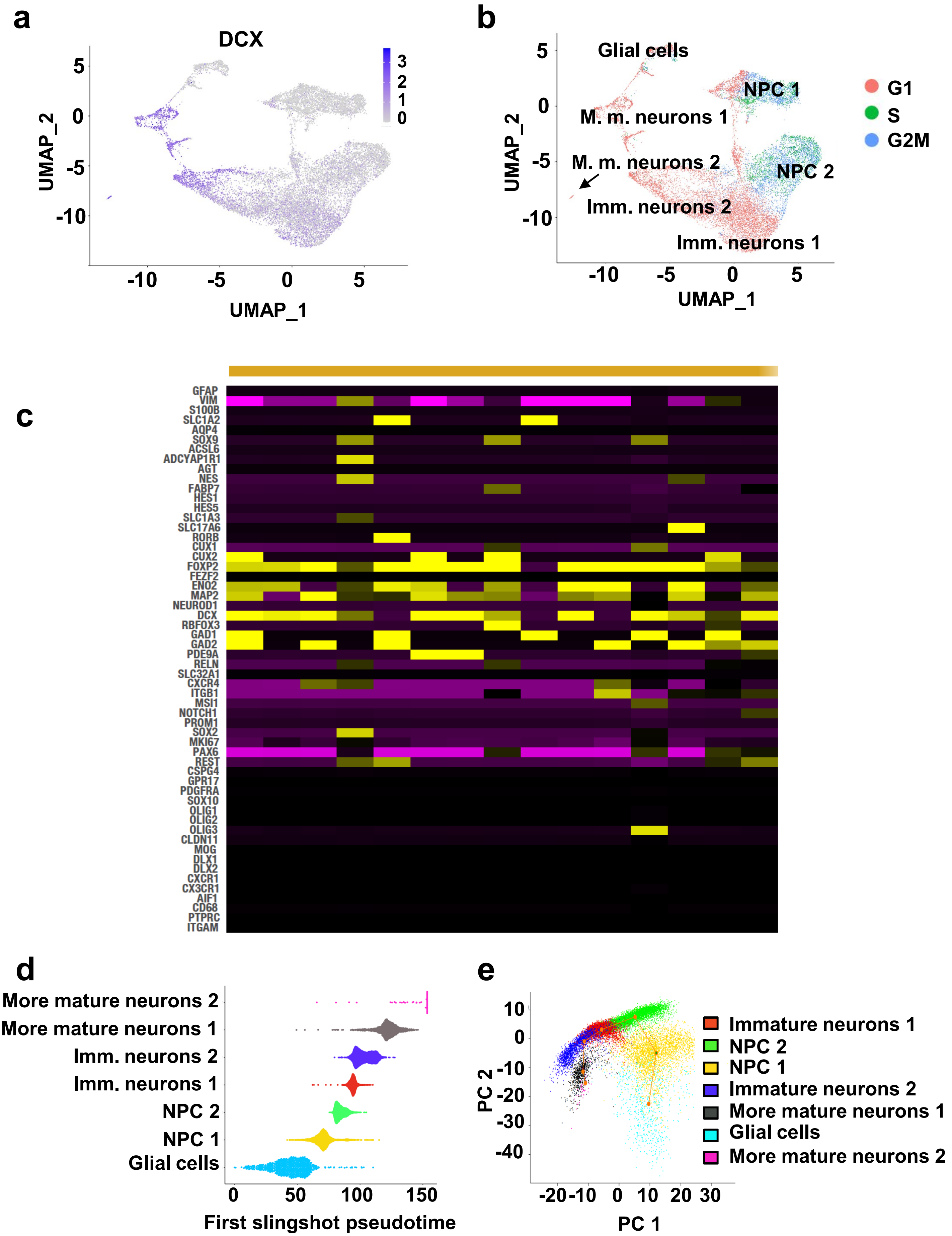

### Suppl. Figure 8

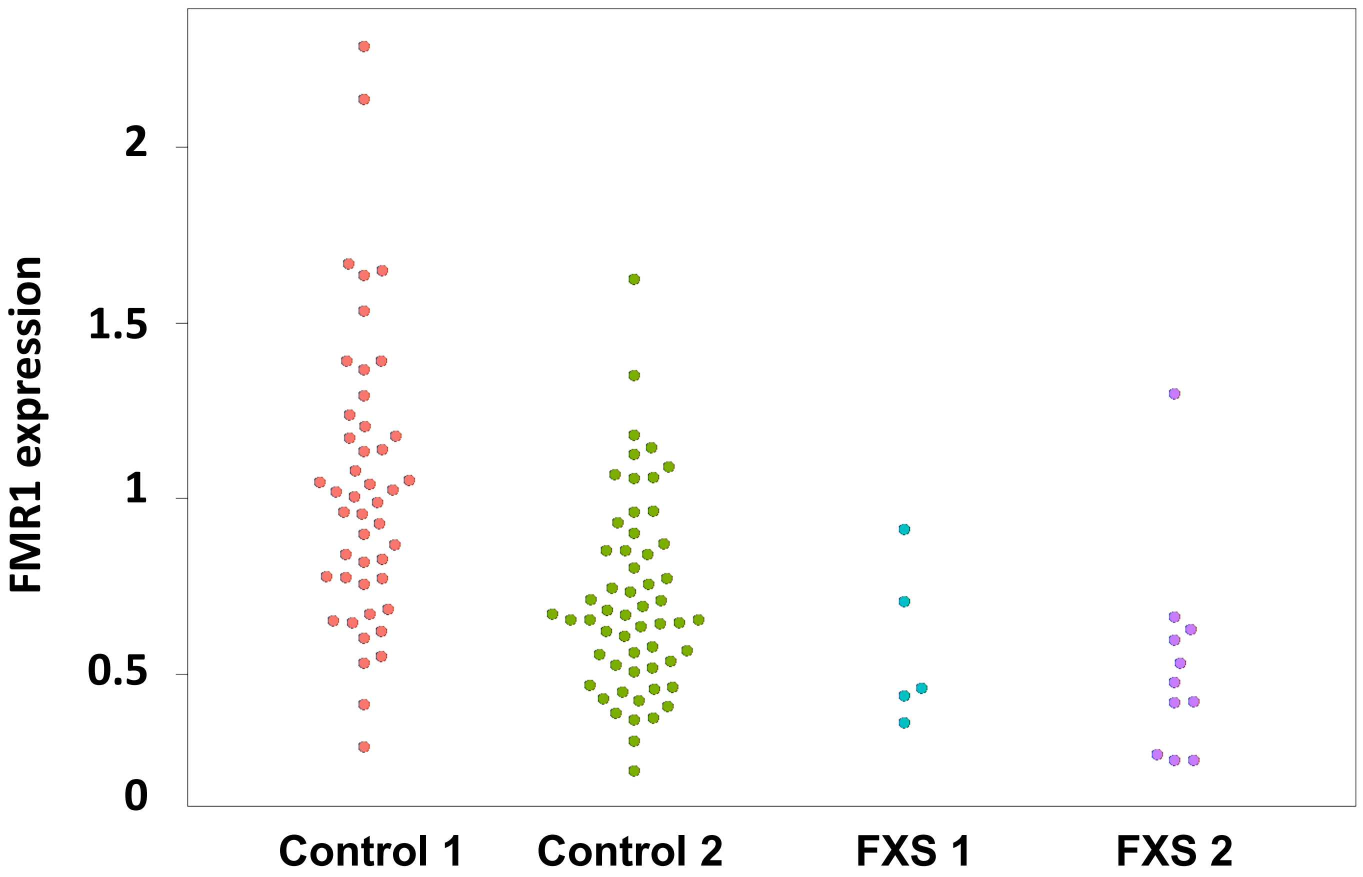

### Suppl. Figure 9

**a**

| Cluster | Control 1 | Control 2 | FXS 1 | FXS 2 | FC | p-value (BH) |
|---|---|---|---|---|---|---|
| NPC 1 | 830.9 | 945.0 | 732.6 | 770.5 | -1.2 | 0.03 |
| NPC 2 | 1064.3 | 933.9 | 739.9 | 815.6 | -1.3 | 0.06 |
| Immature Neurons 1 | 1150.0 | 1243.5 | 901.1 | 914.7 | -1.3 | 0.02 |
| Immature Neurons 2 | 645.9 | 661.4 | 802.2 | 910.2 | 1.3 | 0.02 |
| More mature Neurons 1 | 206.0 | 122.6 | 395.6 | 319.9 | 2.2 | 0.02 |
| More mature Neurons 2 | 5.7 | 1.2 | 80.6 | 60.8 | 17.4 | 0.0003 |

**b** NPC

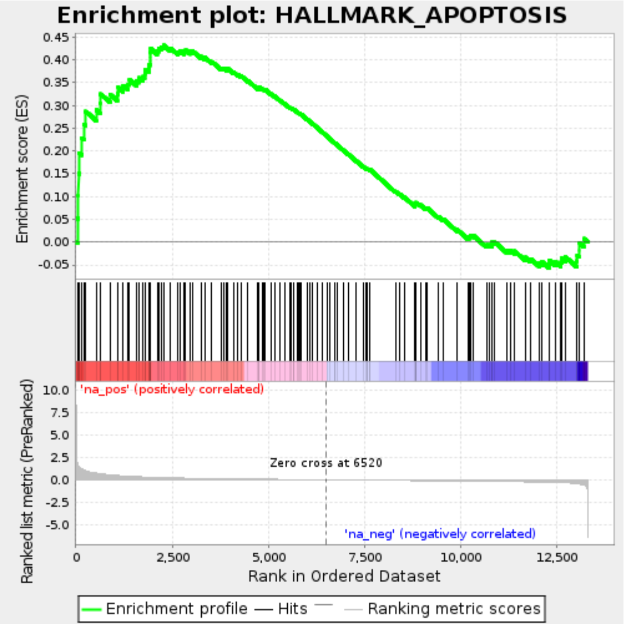


**c** Neurons

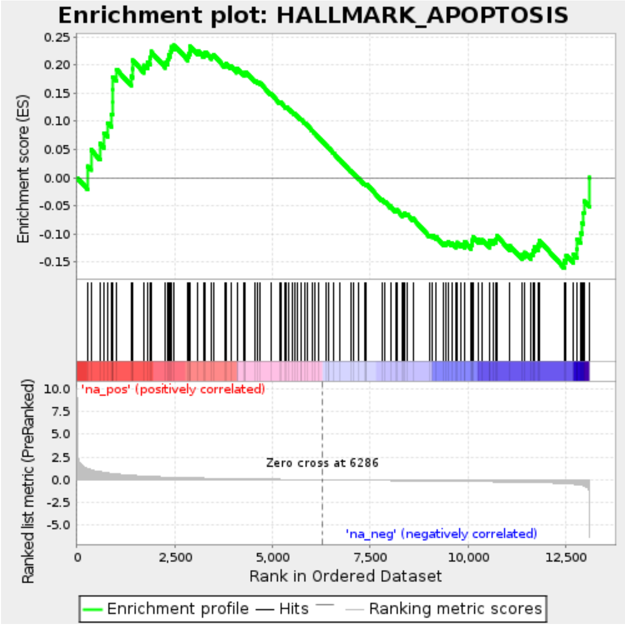


**d**

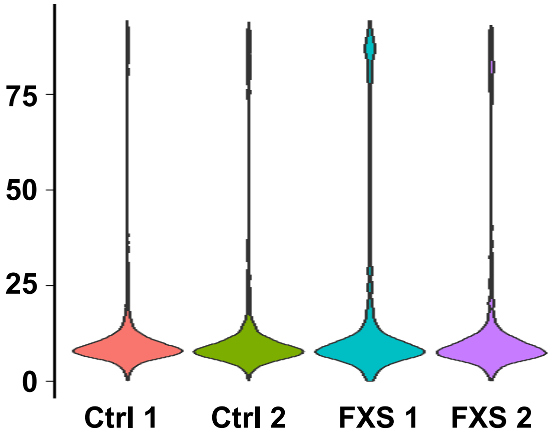

### Suppl. Figure 10

| Gene | $log_2FC$ | P-value |
|---|---|---|
| **ARC** | 1.3 | 0.007 |
| **FOS** | 0.3 | 0.01 |
| **EGR1** | 0.3 | 0.03 |

.

### Suppl. Figure 11

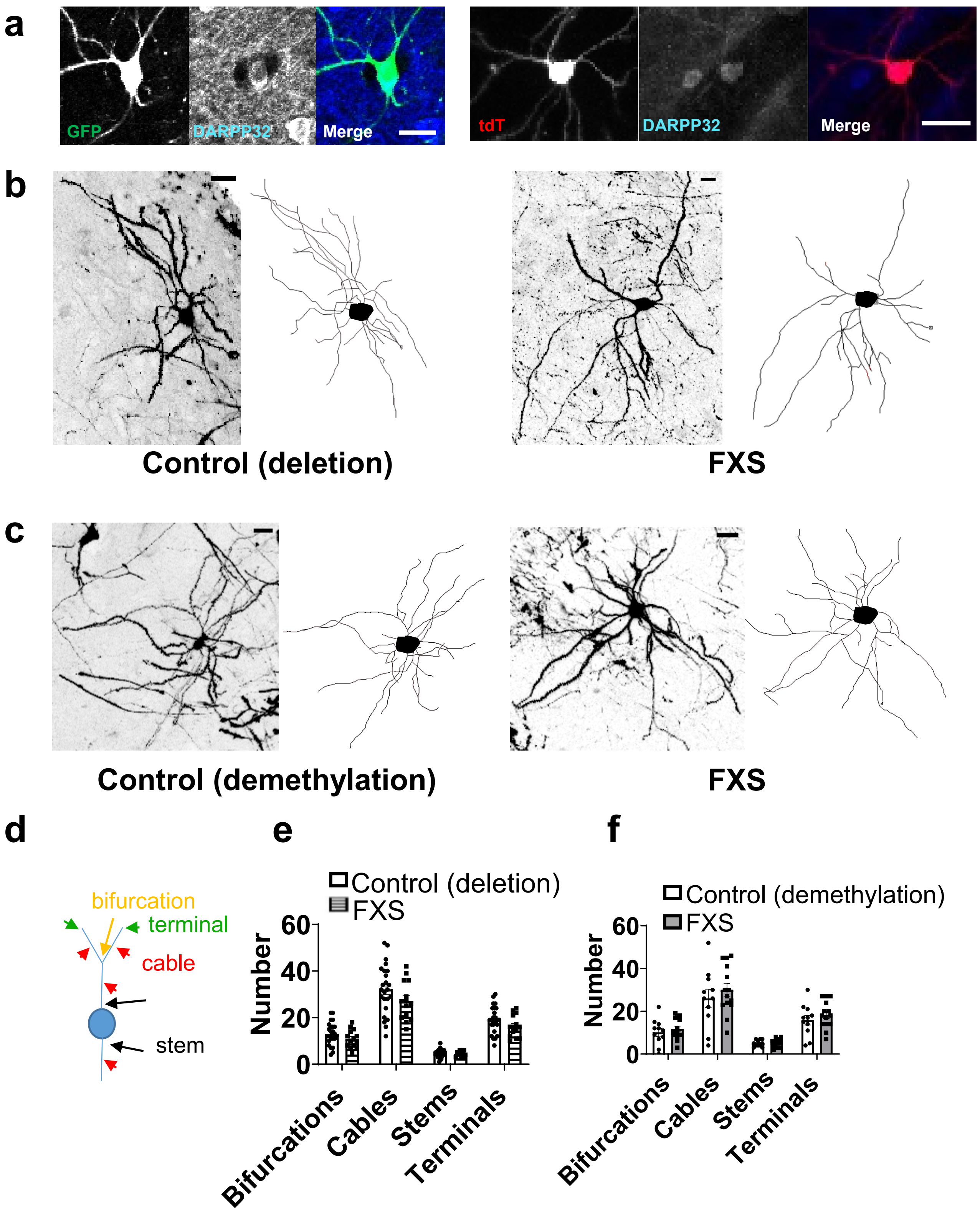
